# Privacy-preserving quality control of neuroimaging datasets in federated environment

**DOI:** 10.1101/826974

**Authors:** D. K. Saha, V. D. Calhoun, Y. Du, Z. Fu, S. R. Panta, S. Kwon, A. D. Sarwate, S. M. Plis

## Abstract

Privacy concerns for rare disease data, institutional or IRB policies, access to local computational or storage resources or download capabilities are among the reasons that may preclude analyses that pool data to a single site. A growing number of multi-site projects and consortia were formed to function in the federated environment to conduct productive research under constraints of this kind. In this scenario, a quality control tool that visualizes decentralized data in its entirety via global aggregation of local computations is especially important, as it would allow the screening of samples that cannot be jointly evaluated otherwise. To solve this issue, we present two algorithms: decentralized data stochastic neighbor embedding, dSNE, and its differentially private counterpart, DP-dSNE. We leverage publicly available datasets to simultaneously map data samples located at different sites according to their similarities. Even though the data never leaves the individual sites, dSNE does not provide any formal privacy guarantees. To overcome that, we rely on differential privacy: a formal mathematical guarantee that protects individuals from being identified as contributors to a dataset. We implement DP-dSNE with AdaCliP, a method recently proposed to add less noise to the gradients per iteration. We introduce metrics for measuring the embedding quality and validate our algorithms on these metrics against their centralized counterpart on two toy datasets. Our validation on six multi-site neuroimaging datasets shows promising results for the quality control tasks of visualization and outlier detection, highlighting the potential of our private, decentralized visualization approach.

## 1. Introduction

Even though the availability of public data continues to increase, there are still many “unsharable”, private datasets which arise multiple challenges for machine learning systems [1]. The importance of operating on decentralized sensitive data and, as a result, of (virtually) pooling very large-scale neuroimaging datasets is exemplified by the success of the ENIGMA project [2]. The continuing growth in the value of large and diverse neuroimaging datasets should inevitably increase the demand for similar decentralized consortia. Multiple large consortia, such as the Global Imaging Genetics in Adolescents (GIGA) consortium [3], are already leveraging decentralized approaches. Several decentralized systems are being developed to virtually pool and facilitate computation on distributed datasets, for example COINSTAC [4] and others [5, 6, 7, 8]. For all of them, quality control is essential.

Intuitive visualization of the complete virtual dataset that is physically spread across multiple locations is a much-needed tool for filtering out participating sites with bad data, detecting incorrect processing, or identifying mistakes in the input process. For example, consider a magnetic resonance image (MRI) data sample that consists of the entire brain, containing on the order of 100,000 volumetric pixels (voxels) [9]. Outliers in smaller datasets at consortium sites make statistical analyses of the consortium data much more difficult. One solution is to develop methods for quality control of large-scale brain imaging data. Since it is challenging to scan through each data sample, an effective method of quality control is to simultaneously embed multiple samples onto a lower dimensional space for visualization. These visualizations have been shown to be useful tools to assess and monitor data quality, while revealing interesting relationships [10]. Beyond data quality, we can also use these approaches to visualize relationships among groups (e.g. diagnostic categories) or continuous measures (such as disease severity or cognitive performance) [11].

A common way of visualizing a dataset consisting of multiple high dimensional data points is to embed them into a 2 or 3-dimensional space. Existing methods like principal component analysis (PCA) [12] can be useful for revealing the linear structure of data. However, the nonlinearity of biomedical data makes analysis with PCA difficult, failing to preserve and convey the hidden structure within the data. To resolve this issue, many other methods, including Sammon mapping [13], curvilinear component analysis [14], stochastic neighbor embedding [15], isomap [16], maximum variance unfolding [17], locally linear embedding [18], and Laplacian Eigenmaps [19] were developed to embed and visualize nonlinear datasets. These methods perform well on artificial data, but can struggle in real high dimensional settings due to their inability to retain local and global structure in a single map. Several methods have been proposed to overcome these problems as well. To visualize underlying structure and intrinsic transitions in high-dimensional biological data, an approach that is highly scalable both in memory and runtime, called potential of heat diffusion for affinity-based transition embedding (PHATE), was recently introduced [20]. Other notable methods include t-distributed stochastic neighbor embedding (t-SNE) [21], viSNE [22], and hierarchical stochastic neighbor embedding (HSNE) [23]. Lastly, to reduce dimensions and overcome computational restrictions, Uniform Manifold Approximation and Projection (UMAP) [24] was proposed and has proven to be effective in the field of bioinformatics. However, all of these methods were built on the principle that the datasets are locally accessible. If the data samples were distributed across multiple sites, the sites would have to pool their data to a single site for analysis.

In this paper, we propose decentralized stochastic neighbor embedding (dSNE), an algorithm that embeds a high dimensional, decentralized dataset into a 2D map for subsequent visualization and inspection. Our approach improves on our preliminary adopts the method of embedding multiple modalities into the same Euclidean space based on their co-occurrence statistics [25]. Since we cannot physically pool all of the data to a single local site, we use publicly available anonymized datasets as a common reference and build the overall embedding around it. This approach is most similar to the method of landmark points, previously used for improving computational ef-ficiency [26, 27]. Our approach, dSNE, significantly extends the original landmark points approach, using t-SNE as the base of our algorithm. Our method can be seen as a dynamic modification that can embed data points into a common space after capturing the relationship among samples distributed across different locations. It significantly improves on our prior work [28] by using multiple iterations to improve the embedding. Even though dSNE provides a way to visualize data in a decentralized manner, it comes with no formal privacy guarantees. To remedy this, we also propose an (*ϵ*, *δ*)-differentially private version of dSNE (DP-dSNE). Differential privacy is a framework which quantifies the privacy risk to individuals when functions of their data are released to untrusted parties. DP-dSNE adds noise to the gradients of the private and shared data per iteration, using a method called AdaCliP. We evaluate our DP-dSNE algorithm using Rényi Differential Privacy [29] and present a privacy analysis using the moments accountant to keep track of the privacy loss per iteration. To evaluate and compare the performance with the centralized version in controlled settings, we demonstrate both of our algorithms (dSNE and (*ϵ*, *δ*)-DP dSNE) on multiple datasets. We also introduce a novel performance metric of overlap and roundness to quantify the quality of our embeddings. Lastly, we apply our approaches to six different multi-site neuroimaging datasets, showing that our methods can capture information and perform quality control of distributed datasets producing highly pragmatic visualizations.

## 2. Methods

In the centralized problem of data embedding, we are given the task of producing a dataset of *N* samples **Y** = [**y**_1_…, **y**_*N*_], where 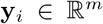, from a dataset **X** = [**x**_1_…, **x**_*N*_], where 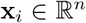, such that *m* ≪ *n*. For the convenience of visualization, *m* is usually set to *m* = 2. The goal is for the embedding **Y** to give a “faithful” embedding of the data **X** in the sense that similar points in **X** will be mapped to close points in **Y**.

### 2.1. Background: t-SNE

In t-SNE, the distances between the points in **Y** must be as close to the distances between the corresponding points in **X**, where preserving the closeness of nearby points is weighted more heavily than far points [21]. In the first step, t-SNE converts the high dimensional Euclidean distances between datapoints into conditional probabilities, called pairwise affinities, that represent similarities between data points (see Algorithm 1). The algorithm takes a scalar parameter called the *perplexity ρ*. To compute similarity of a datapoint *x_j_* to datapoint *x_i_*, the algorithm first computes the weight of *x_j_* given by a Gaussian kernel centered at *x_i_* with bandwidth (variance) *σ_i_*(*ρ*)^2^, where we identify the value of *σ_i_* separately for each datapoint by performing a binary search across a range of values until we can match the user-specified perplexity. The similarity is a conditional probability distribution *p_j|i_* formed by renormalizing the *N* likelihoods into a probability mass function:

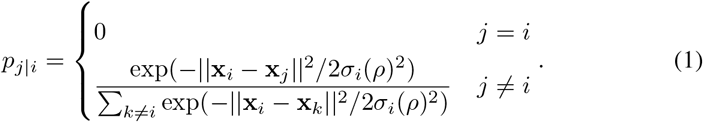

These similarities can be gathered into an *N* × *N* matrix 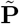 and the affinity is the symmetrized matrix 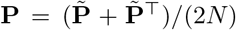. This makes **P** a matrix representing a joint distribution on pairs of data points.

**Algorithm 1.**
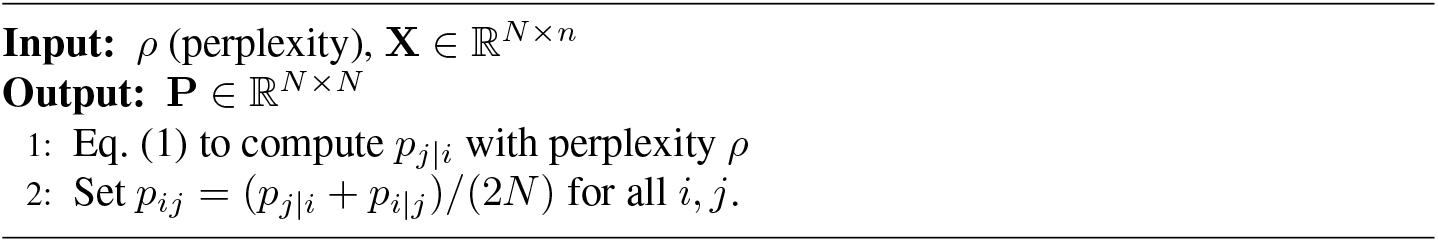
PairwiseAffinities

For the low-dimensional representation **Y** we compute pairwise weights in a similar way, except that this time for the joint distribution *q_ij_* we use the Student-*t* distribution with one degree of freedom (or a Cauchy distribution) instead of a Gaussian:

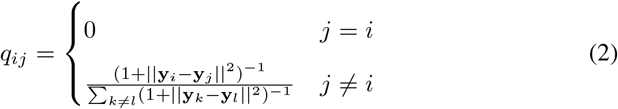

These can be gathered into a matrix **Q** representing a joint distribution for the embedded points. Algorithm 2 outlines the full t-SNE procedure. To embed the data points into a low dimensional space, t-SNE tries to minimize the mismatch between distribution **P** and **Q** in higher and lower dimensional spaces. The algorithm performs gradient descent on the Kullback-Leibler (KL) divergence (or relative entropy) between the joint distribution **P** and the joint distribution **Q**:

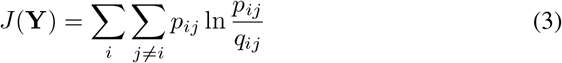

The gradient of the Kullback-Leibler divergence between **P** and the Student-t based joint probability distribution **Q** is expressed in equation (4).

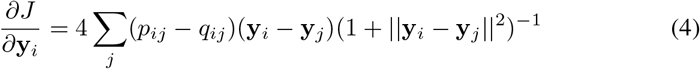

**Algorithm 2.**
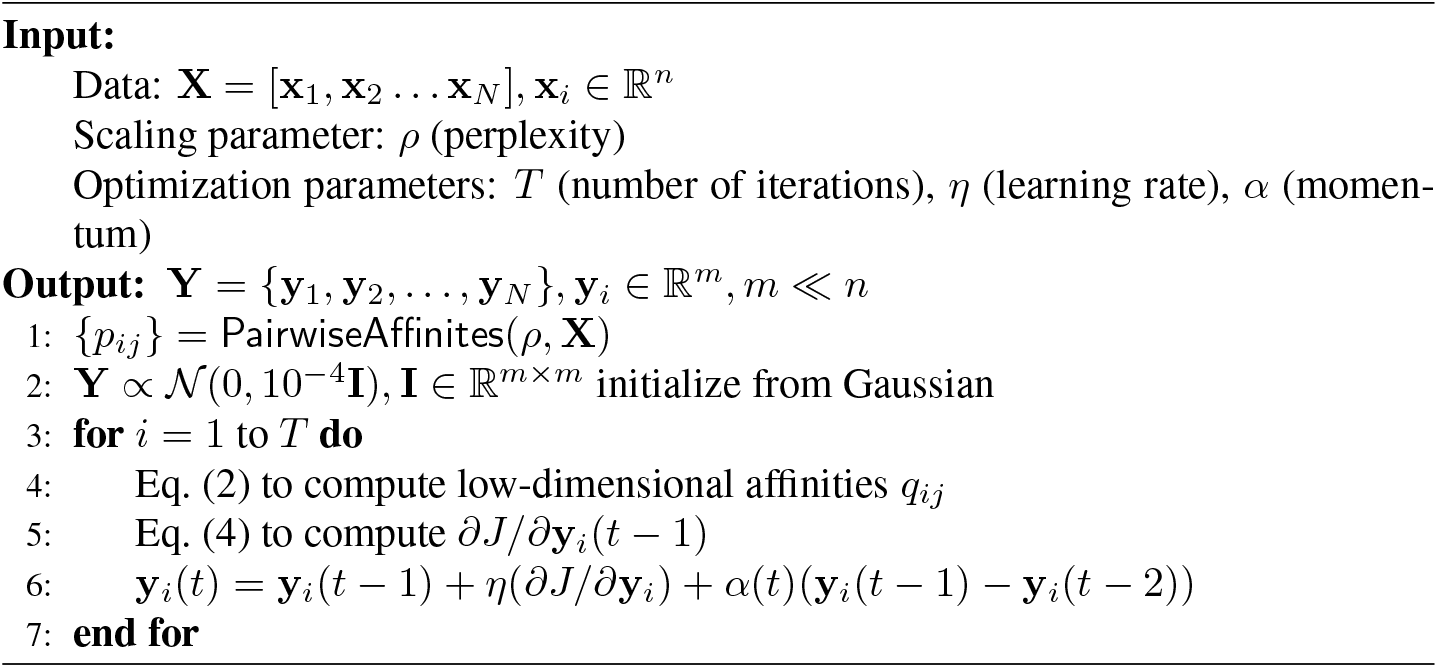
tSNE

Inspired by the overall satisfactory performance of t-SNE on a range of tasks, we use it as the base of our decentralized algorithm.

### 2.2. Proposed method: dSNE

In the decentralized setting, the privacy and sensitivity of datasets often preclude the pooling of local data, making computation of distances amongst samples across different sites difficult. Without these distances (see equation (1)), we cannot obtain a common embedding. Fortunately, in neuroimaging (and many other fields), there are now multiple large public repositories of MRI data that we can leverage to make this computation feasible [30, 31, 32].

In the decentralized setting we have *L* sites where each site ℓ has (local) data 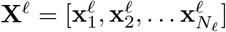 consisting of *N*_ℓ_ vectors **x**^ℓ^ in 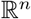. In addition we have a shared data set 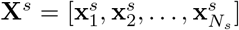. The goal is to produce embeddings {**Y**^ℓ^} and **Y**^*s*^, where 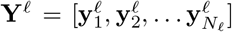 for each ℓ and 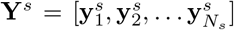 contain vectors in 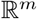 where *m* ≪ *n*. Typically we will consider *m* = 2 to produce 2D visualizations of the data set. As in t-SNE, we want the distances close **x** points to remain close in the embedding of **y** points. We assume all sites have access to the shared data **X**^*s*^ and its embedding **Y**^*s*^ and can modify **Y**^*s*^ when they update locally.

We implemented three algorithms: (1) Single-shot dSNE, (2) Multi-shot dSNE, and (3) Differentially private multi-shot dSNE, which we describe in the followiing subsections. Detailed procedure and experimental results of Single-shot dSNE are provided in our prior work [28] and in Appendix A.

#### 2.2.1. Multi-shot dSNE

For multi-shot dSNE, we pass messages iteratively between the local sites and central site in rounds. At time *t*, the centralized site passes the reference embedding **Y**^*s*^(*t* – 1) from the previous iteration to each of the local sites. At this point each site ℓ has **X**^ℓ^, past values of **Y**^ℓ^(*t* – *j*) for *j* = 1,2,…, *t*, the reference data set **X**^*s*^, and the updated embedding **Y**^*s*^ (*t* – 1). Each local site then computes the gradient update (Algorithm 3) using a “momentum” approach that combines information from the past two iterations. The result are updates **Y**^ℓ^(*t*) and **Y**^*s*,ℓ^(*t*) for the local and shared data embeddings respectively. The sites send their new embeddings of the local data **Y**^*s*,ℓ^(*t*) to the central site, which averages them to form 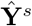 and sends that back to the local sites.

**Algorithm 3.**
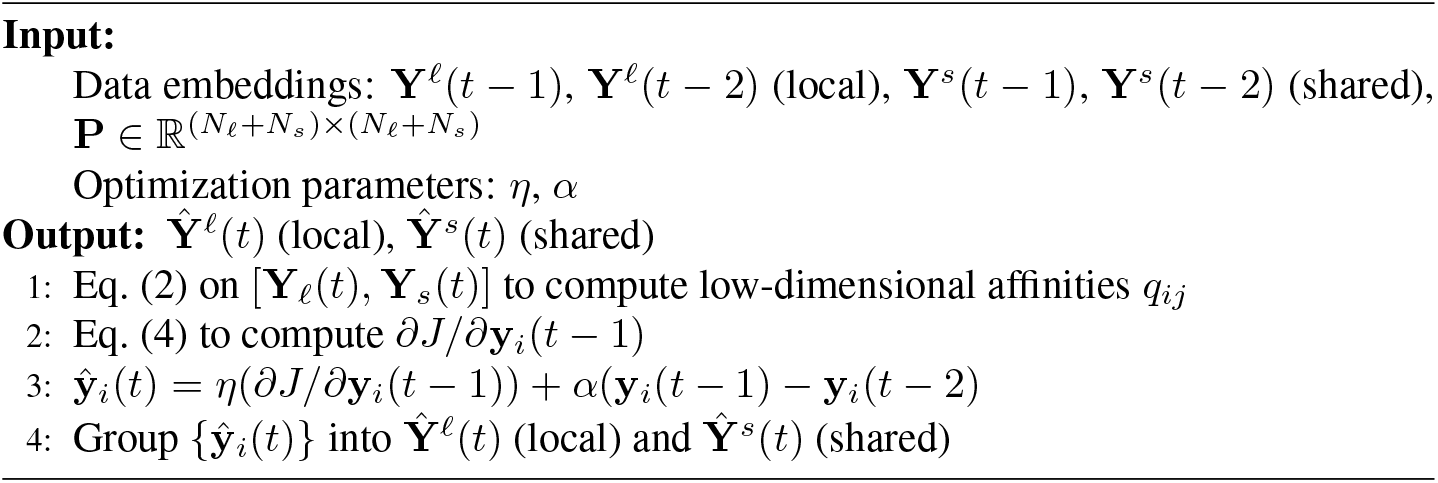
LocalGradStep

**Algorithm 4.**
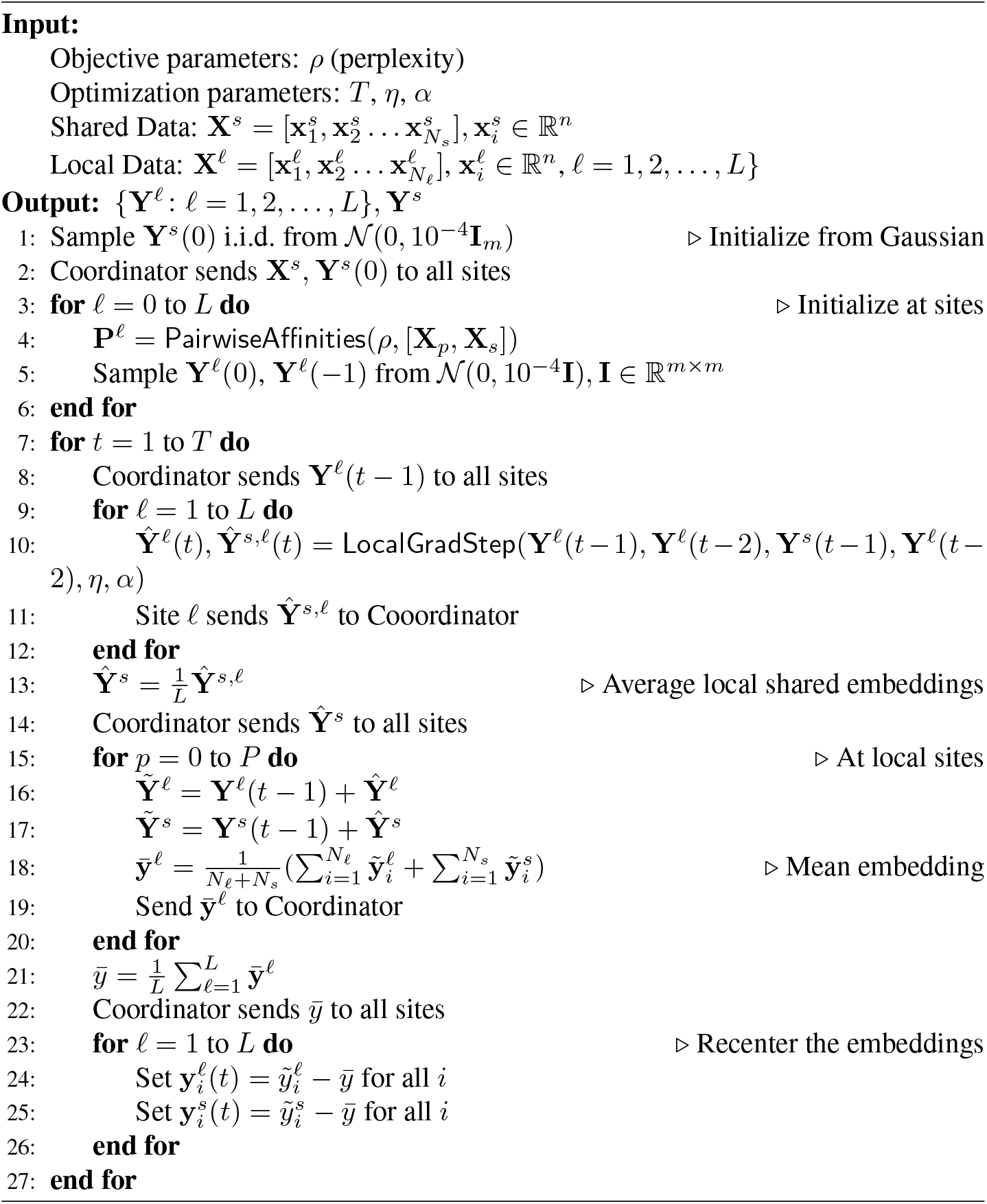
multishotDSNE

The local sites then update their local and shared embeddings and compute the average of all embedding to help recenter. They send this average to the coordinator, who averages across sites and sends back a global mean that sites use to center their local and shared embeddings to get **Y**^(ℓ)^(*t*) and **Y**^(*s*)^(*t*) for the next iteration. The pseudocode and overall procedure for multi-shot dSNE are shown in Algorithm 4. Note, that at each iteration, the embedded vector *Y* for the shared dataset will be the same at all of the local sites. This ensures that the local values of different sites are influenced by the same and common reference data at each iteration.

### 2.3. Differentially Private Multi-shot dSNE

We begin by reiterating the setup of multi-shot dSNE. In dSNE, there are n local sites (each with their own disjoint dataset) that would like to collaborate to learn a global interrelational structure among the data samples. However, the sensitivity of biomedical data prevent centralized analyses that pool all the data to a single site. Even though the local data samples never leave the sites in dSNE, since the embeddings of the shared data are influenced by the local data points, there is room for a potential privacy breach. To resolve this, we introduce DP-dSNE, a differentially private dSNE algorithm that formally guarantees privacy. We now define differential privacy [33] and the AdaCliP algorithm [34].

We say that two datasets *D*, *D*′ are *neighboring* datasets if they differ by one data entry. A randomized mechanism 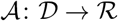 is said to be (*ϵ*, *δ*)-differentially private if for all neighboring databases 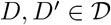, and any measurable set 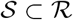, we have

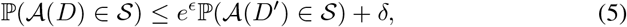

where 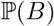 is the probability of the event *B* and the probability is taken over the randomness in the mechanism 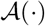.

We use the shorthand notation (*ϵ*, *δ*)-DP for (*ϵ*, *δ*)-differentially private. A standard method to preserve privacy of a function is to add noise, where the variance of the noise is proportional to the sensitivity of the function. Mathematically, we define the ℓ_2_ global sensitivity of a function *f* as

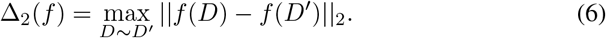

**Algorithm 5.**
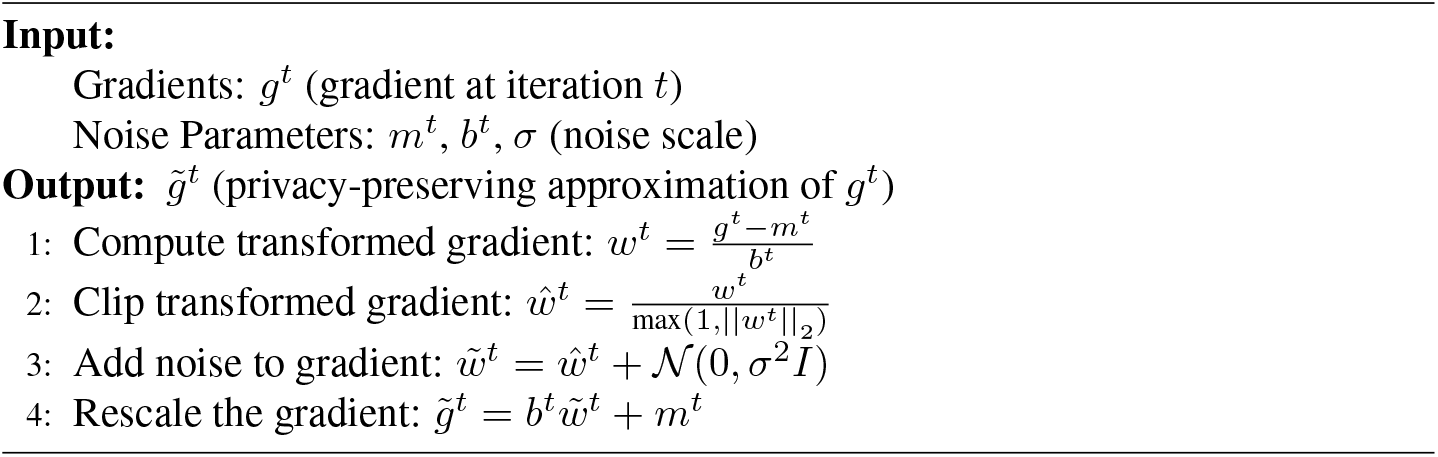
noiseAddition

**Algorithm 6.**
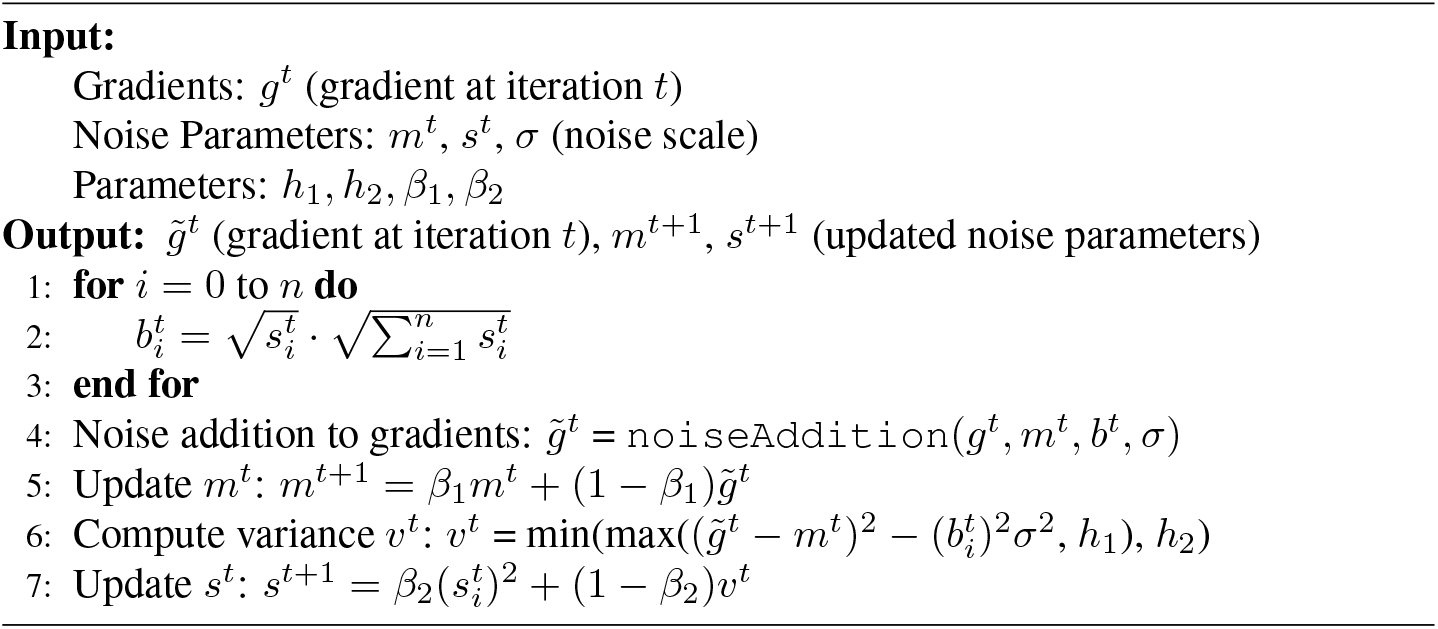
AdaCliP

Given that the ℓ_2_ sensitivity of a function *f* is Δ, one way to preserve privacy is to add Gaussian noise [35] of variance Δ^2^*σ*^2^, such that 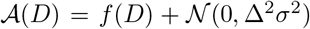. Then, if we choose *σ* to be 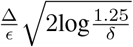, each iteration of the algorithm is (*ϵ*, *δ*)-DP [35], [36]. However, in practice, finding an a *priori* bound on the size of the gradients (i.e. the sensitivity of the gradients) is difficult, and often does not exist. In literature, one way to solve this issue is to bound the gradients by clipping each gradient in ℓ_2_ norm for a clipping threshold *C* [36]. This clipping would ensure that the sensitivity of the gradients change by at most *C*. Although this is a plausible method, clipping all of the gradients to a fixed value of *C* can often add more noise than needed, as the size of the gradients generally grow smaller during training.

To add less noise per iteration, we adopt in using AdaCliP [34], a recently proposed method to adaptively clip and add noise based on the size of the gradients. By introducing AdaCliP into our multi-shot dSNE algorithm, we can preserve the privacy of the individuals in the dataset. The modification that we need in order to make dSNE (*ϵ*, *δ*)-DP is to replace `LocalGradStep` in Algorithm 4 with `DP-LocalGradStep`, as shown in Algorithm 7. There are two things to note about the DP algorithm: (1) `DP-LocalGradStep` brings two extra parameters, *m* and *s* that we use and update at every iteration and (2) even though we are computing a global mean based on each local site’s mean, each step is still differentially private due to post-processing invariance [35]. In the next section, we provide a privacy analysis using Rényi Differential Privacy and use the moments accountant to keep track of the total privacy loss during training [36].

**Algorithm 7.**
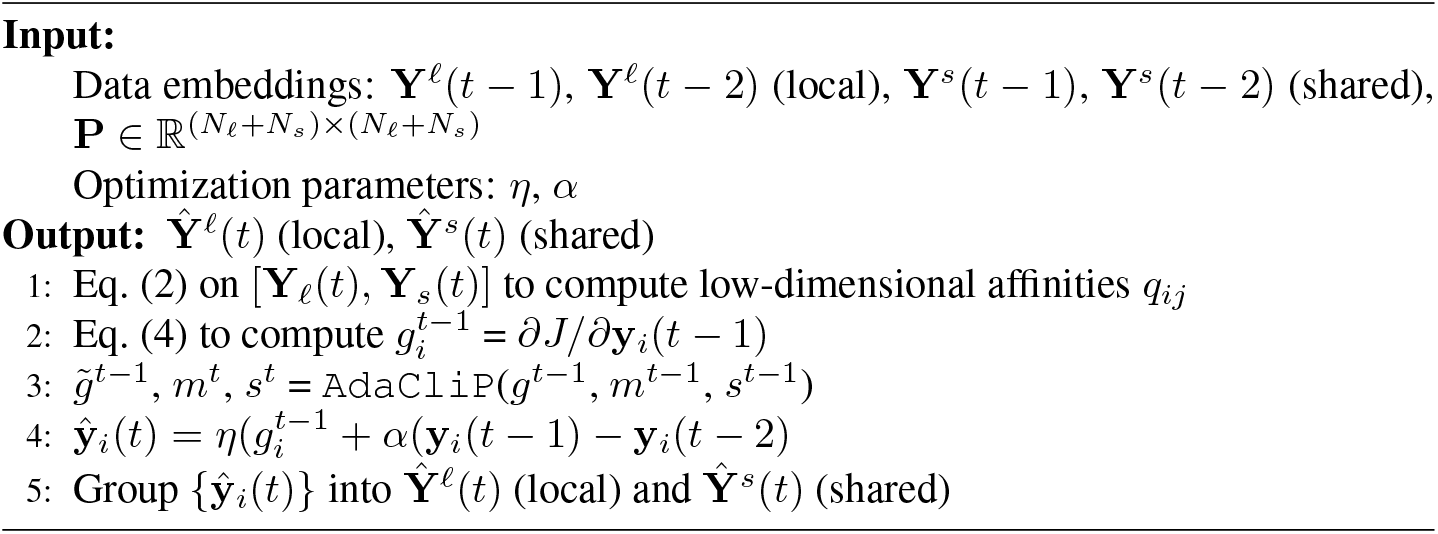
DP – LocalGradStep

### 2.4. Privacy Analysis of Differentially Private Multi-shot dSNE

In this section, we analyze the privacy loss of our differentially private multi-shot dSNE algorithm using Rényi Differential Privacy (RDP) [29]. Recall that in our decentralized setup, each local site has data that is disjoint from the other local sites. Thus, it is sufficient to analyze the privacy loss random variable of a single site. We start our analysis of the DP-dSNE algorithm by reviewing some definitions and connections between RDP and DP.

If 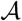 is a randomized algorithm satisfying (*α*, *γ*)-RDP, then it also satisfies 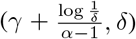-DP for any 0 < *δ* < 1. Further, if 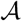 has ℓ_2_ sensitivity Δ, then the Gaussian mechanism 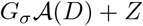, where 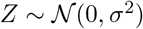, satisfies 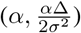-RDP [29]. We can now derive the sensitivity of our gradients from using AdaCliP to define an (*ϵ*, *δ*)-DP guarantee.

Let 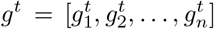 and 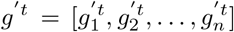 be gradient vectors of two neighboring datasets at iteration *t*. Using AdaCliP, we have two additional vectors for each dataset: 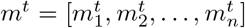 and 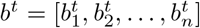. The sensitivity of the gradients of the neighboring databases yield

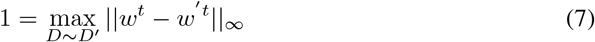

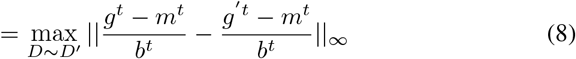

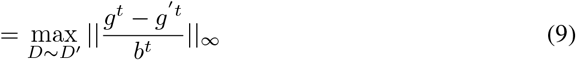

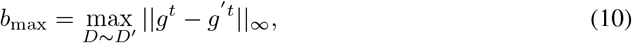

where ‖**x**‖_∞_ = max_*i*_ |*x_i_*| (i.e. the ℓ_∞_ norm a vector **x** with *i* entries) and *b*_max_ = ‖*b^t^*‖_∞_. Since the transformed gradient *w^t^* is clipped at norm 1, its sensitivity is also 1. Following this analysis, we obtain that the sensitivity of the gradient *g^t^* is *b*_max_.

By using the definitions stated previously, if *J* denotes the number of iterations, the DP-dSNE algorithm satisfies 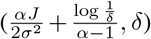-DP, where *σ*^2^ is the variance of the noise, 0 < *δ* < 1, and *α* is

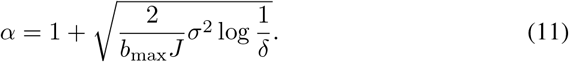

At each iteration, we keep track of *b*_max_ to compute our (*ϵ*, *δ*)-DP guarantee. Note that in our privacy analysis, we defined our (*ϵ*, *δ*)-DP guarantee in terms of RDP, but we can easily derive these values using the moments accountant [36], similar to that of [37].

### 2.5. Comparison Metrics

To measure the performance and quality of clustering, several metrics have been proposed, including the Davies-Bouldin (DB) index [38], Dunn index [39], quality index [40], Bayesian information criterion (BIC) index [41], and the silhouette coefficient [42]. Other metrics such as F-measure, entropy, purity, and rand index can also be used as an external comparison metric.

In the distributed setting, it is important to make sure that the embedding of a data point gets clustered into its correct corresponding class. To test the cluster qualities of our decentralized algorithms, we use two a priori labeled datasets: (1) the MNIST dataset and (2) the COIL-20 dataset, whose clusters are known and described [21]. We consider each label as the known number of clusters as our ground truth. We introduce three new validation techniques: (1) K-means ratio (2) Intersection area and (3) Roundness.

1. The k-means criterion (or ratio) is the ratio of intra and inter-cluster distances between clusters. Mathematically, this is defined as

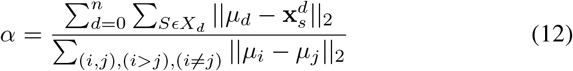

where *n* is the number of clusters. We can interpret this ratio as the following: the numerator indicates intra cluster distance, in which a smaller value shows that each distinct cluster is tightly bounded. A larger denominator value implies that the inter clusters are well separated from each other. Thus, if the *α* value is generally small, we can conclude that the algorithm performs well.
2. The intersection area can be seen as computing the overlap between two clusters. We first remove the outliers (see Figure 1) [43] to compute the convex hull for each group and use them to measure the overlap. In specific, this is done by summing all of the polytope areas minus the area of the union of all the polytopes, normalized by the area of the union. The lower value of intersection area determines the formation of good cluster with less overlap.
3. Roundness is the ratio of the area of each polytope to the area of the circumscribed circle. To compute the roundness of each cluster, we first represent each cluster as a cloud with a convex hull. To remove the effect of the differences in perimeters, we normalize the perimeters. This normalization effectively approximates the process of making the perimeters equal. Finally, we compute the area of the polytope and use the area as our measure of roundness. The higher roundness metric value determines the good quality of the cluster. More precisely, it quantifies how the samples are distributed around the mean of a given cluster.

**Figure 1:**
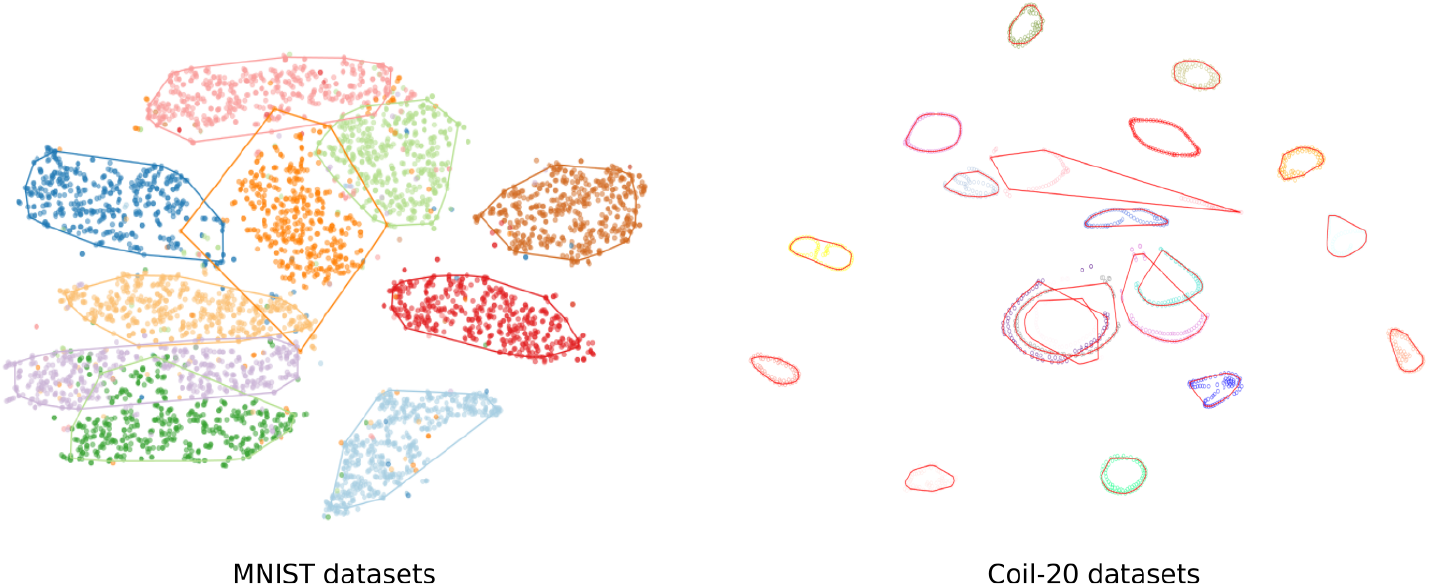
A t-SNE output on centralized MNIST and COIL-20 dataset and outlier-free convex hull boundaries.

## 3. Data

We base our experiments on seven datasets:

1. MNIST dataset^1^ [44]
2. COIL-20 dataset^2^ [45]
3. Autism Brain Imaging Data Exchange (ABIDE) fMRI dataset^3^ [46]
4. Pediatric Imaging Neurocognition Genetics (PING) dataset^4^ [47]
5. Structural Magnetic Resonance Imaging (sMRI) dataset
6. Function Biomedical Informatics Research Network (fBIRN) structural MRI dataset [48] and
7. Bipolar and Schizophrenia Network for Intermediate Phenotypes (BSNIP) structural MRI dataset [48]
8. Functional Magnetic Resonance Imaging (fMRI) dataset

**MNIST** dataset was taken from a Kaggle competition which had 28000, 28 × 28 gray-scale images of all 10 handwritten digits. For our experiments, we randomly chose 5000 different samples from the dataset (while preserving class balance). For centralized t-SNE and dSNE, we pre-processed the dataset by reducing the dimensions of the data samples from 784 to 50 using Principal Component Analysis (PCA).

**COIL-20** dataset contains images of 32 × 32 = 1024 pixels of 20 different objects. Each object was placed on a motorized turntable against a black background. Between 0° and 360° degrees, a picture was taken in 5° degree intervals. Each object was viewed from 72 equally spaced orientations, yielding a total of 1440 images.

**ABIDE** fMRI dataset contains data samples of 1153 subjects accessible through the COINS data exchange (http://coins.mrn.org/dx). The ABIDE dataset was pre-processed down to multiple spatial and temporal quality control (QC) measures^5^. Since this dataset inherently had lower dimensions, we ran t-SNE and dSNE directly on the dataset without a dimensionality reduction step.

**PING** is a multi-site study containing neural developmental histories, information about developing mental and emotional functions, multimodal brain imaging data, and genotypes for well over 1000 children and adolescents between the ages of 3 to 20. We take 632 subject’s fMRI data from this dataset for our experiment. The data is pre-processed with SPM5 [49] software. It is slice time corrected and warped to the standard MNI brain template from SPM5. This image is used for extracting the data for the experiment. For the first time point, the voxel values at each location from all the brain slices are first added across slices along the Z axis, resulting in a single row vector of size 3339. This was done to reduce the computational load on the system, and hence improve run time of the proposed algorithm. These voxel values from each image scan served as inputs to the t-SNE and dSNE algorithms.

**sMRI** scans (3D T1-weighted pulse sequences) are pre-processed through the voxel based morphometry (VBM) pipeline using the SPM5 software. VBM is a technique using MRI that facilitates examination of focal differences in brain anatomy, using the statistical approach of parametric mapping. Gray matter maps are extracted from segmenting the T1 weighted nifti images. The unmodulated gray matter concentration images from the VBM pipeline are registered to the SPM template. In some cases, the non-modulated maps are preferred compared to the modulated maps according to existing literature [50]. The data from these unmodulated gray matter normalized images is used for this experiment. To reduce the computational load on the system, and hence improve run time of the proposed algorithm, for each scan, the voxel values at each location from all the brain slices are first added across slices, resulting in a matrix size of 91 × 109. All of the voxel values from each image scan are converted into a single row vector of size 9919 for each data point and passed as inputs to the t-SNE and dSNE algorithms.

**fBIRN** and **BSNIP** datasets used in this study were collected from seven and six imaging sites, respectively. Each subject was selected based on head motion (≤ 3° and ≤ 3*mm*) and functional data providing nearly full brain normalization [51]. These criteria yielded a total of 311 subjects (160 schizophrenia (SZ) patients and 151 healthy control (HC)) for the fBIRN dataset and 419 subjects (181 SZ and 238 HC) for the BSNIP dataset. In this study, the Neuromark pipeline [48] was adopted to extract reliable intrinsic connectivity networks (ICNs) that were replicated across independent datasets.

**fMRI** dataset consists of 3910 subjects of echo planar imaging data by following different protocols across multiple sites and studies. For pre-processing, the statistical parametric mapping (SPM5) toolbox was used for slice time correction, motion correction, and spatial normalization. For each of these images, six quality control (QC) metrics are computed from this data and are given as inputs to t-SNE algorithm. As the QC matrices values were low dimensional, we directly run t-SNE and dSNE without reducing the dimensions.

## 4. Experimental Setup

In this section, we discuss the experiments used to compare centralized t-SNE to dSNE and DP-dSNE. The experiment with the fMRI dataset illustrates how we can use dSNE for outlier detection and quality control. We use the terms “reference” and “shared” data interchangeably throughout the setup. We organize this section into two parts: (1) experiments on t-SNE and dSNE and (2) experiments on t-SNE and DP-dSNE.

### 4.1. Experiments with dSNE

#### 4.1.1. MNIST Data

For the MNIST dataset, we consider a total of four experiments to compare centralized t-SNE to dSNE. Each of these experiments imitiate a possible scenario between the coordinator and local nodes.

##### Experiment 1 (Effect of the sample size)

The objective of this experiment is to investigate the adaptability of our algorithm when there is an imbalance in the number of data samples between the local site and central node. This imitates a plausible phenomenon, where the centralized site accumulates and stores more data than the local sites (and perhaps, vice versa). In this experiment, each local site holds samples according only one digit (label), while the reference dataset contains all of the digits (0-9). We consider 2 cases: (1) each site contains 400 samples while each digit in the reference consists of 100 samples and (2) the inverse case, in which the local sites have only 100 samples, while the reference has 400 samples.

##### Experiment 2 (No diversity in the reference data)

In this experiment, the reference dataset contains a single digit that is also present at all of the three local sites. We run this experiment for each of the 10 digits. Each site contains a total of 400 samples for each of its corresponding digits, while the reference dataset contains 100 samples of its digit.

##### Experiment 3 (Missing digit in reference data)

In this experiment, we investigate the effect of the case when a digit is missing from shared (reference) data. This approximates the scenario of unique conditions at a local site. Each local site out of 10 contains a single digit. We run a total of 10 experiments, where in each, the reference dataset is missing a digit. For each of the experiments, we have 2 conditions: one in which the reference dataset is small (100 samples for each digit but the missing one) while the sites are large (400 samples per site) and another in which the reference data is large (400 samples per digit) and the sites are small (only 100 samples).

##### Experiment 4 (Effect of the number of sites)

In this experiment, we investigate whether the overall size of the virtual dataset affects the result. Each local site, including the reference dataset, contains all digits (0-9) with 20 samples per digit. We continuously increase the number of sites from 3 to 10. As a result, the total number of samples across all sites increases as well.

#### 4.1.2. COIL-20 Data

##### Experiment 1 (Effect of the sample size)

Similar to the first experiment of MNIST, we apply the same strategy in which the centralized node holds more data than the local sites. In this experiment, every local site contains only one type of object and the reference dataset contains all objects (1 to 20). We consider two cases: (1) each site contains 52 samples of its corresponding object and each object in the reference consists of 20 samples (2) and the inverse case, when each site holds 20 samples, while the reference objects are represented by 52 samples.

##### Experiment 2 (Missing object in reference data)

In this experiment, we investigate the scenario in which some objects are missing from the shared dataset, similar to that of Experiment 3 of MNIST. Each local site out of 20 contains a single object. We run 10 experiments with different random seeds, where in each run, the reference dataset is missing objects from local site 16-20. For each of the experiments, we have 2 conditions: (1) the reference dataset is small (20 samples for each object) while the sites are large (52 samples per site) and (2) the reference data is large (52 samples per object) and the sites are small (only 20 samples).

#### 4.1.3. ABIDE fMRI Data

To simulate a consortium of multiple sites, we randomly partition the ABIDE dataset into ten local and one reference dataset. We run three different dSNE experiments, each corresponding to one random split. We use this dataset to also demonstrate how we can perform quality control of data samples using dSNE. Lastly, we collect all of the data into one local site to simulate a centralized visualization using t-SNE to compare to our dSNE algorithm.

#### 4.1.4. sMRI Data

The sMRI dataset consists of subjects with four different age groups: below 11, 11 to 17, 30 to 34, and above 64. We run a total of three experiments using this dataset:

1. We use each age group as a unique local site (four sites in total) and form reference samples by taking 100 samples from each local site.
2. For this experiment, we keep the local sites the same as the previous setting, but create reference data by taking 100 samples from only the first site.
3. In this case, we randomly take 50 samples from site 2 and place them in site 1 and take 100 samples from site 4 and distribute them equally between sites 2 and 3. This analyzes the effect in which each local site has data samples from the same class.

#### 4.1.5. PING Data

We collect our PING dataset from five different data to run four different experiments:

1. For the first experiment, we use each data source (total of five) as a local site and form our reference samples by taking small samples from each site.
2. The second experiment also uses five local sites, but we form the reference dataset by taking 100 samples from only the second site.
3. Similar to Experiment 3 of the sMRI dataset, we randomly take 30 samples from site 2 to place in site 1, take 20 samples from site 3 to place in site 2, take 10 samples from site 2 to place in site 3, and take 10 samples from site 1 to place in site 4. The reference samples are formed by taking small samples from each local site.
4. For this experiment, we keep sites 1, 3, and 4 unchanged from the first experiment, but form reference samples by taking all of the samples from sites 2 and
5. This yields a total of 3 local sites and one reference sample, consisting of data from sites 2 and 5.

#### 4.1.6. fBIRN and BSNIP Data

For the dSNE experiment, as we collected the fBIRN data from seven different sites, we considered each of them as a local site. The BSNIP dataset (collected from six imaging sites) served as our reference dataset. For t-SNE, we treated each dataset as its own and ran t-SNE separately. We also ran t-SNE on combined datasets (fBIRN + BSNIP) for visual comparisons with dSNE.

#### 4.1.7. fMRI Data

We collected fMRI data from three different sites (MRN, Avanto, and Boulder). For the dSNE experiment, we use each data site as its own unique local site (total of three local sites). However, we randomly picked 200 samples from site 2 and placed them in site 1. Sites 2 and 3 had its own corresponding data from Avanto and Boulder, respectively. We formed the reference dataset by picking 1383 samples from site 1 and 622 samples from site 2. For comparison, we pooled data from all three sites and performed a centralized t-SNE analysis.

### 4.2. Experiments with DP-dSNE

For the DP-dSNE experiment, we used the MNIST and PING datasets to demonstrate the robustness of dSNE even in formal private settings. For the MNIST dataset, we manually created three local sites and one coordinator node to participate in the computation. Each local site holds two classes (digit) from the dataset and the co-ordinator node contains all classes(four in total). For the PING dataset, we used the same setup as stated in Experiment 1 of section 4.1.5. We compare our results to centralized t-SNE and dSNE and show that our algorithm still provides good utility. We hypothesize that our DP-dSNE algorithm will generalize well to other neuroimaging data.

## 5. Results

We organize this section into three parts: (1) comparison between t-SNE and dSNE, (2) comparison between t-SNE and dSNE on biomedical data (3) comparison between t-SNE and DP-dSNE.

### 5.1. Comparison Between t-SNE and dSNE

#### 5.1.1. MNIST Data

For the MNIST experiments, we only present the results of Experiment 1. The other experimental results (Experiments 2, 3 and 4) are presented in [28]. Figure 2 represents the results of Experiment 1.

**Figure 2:**
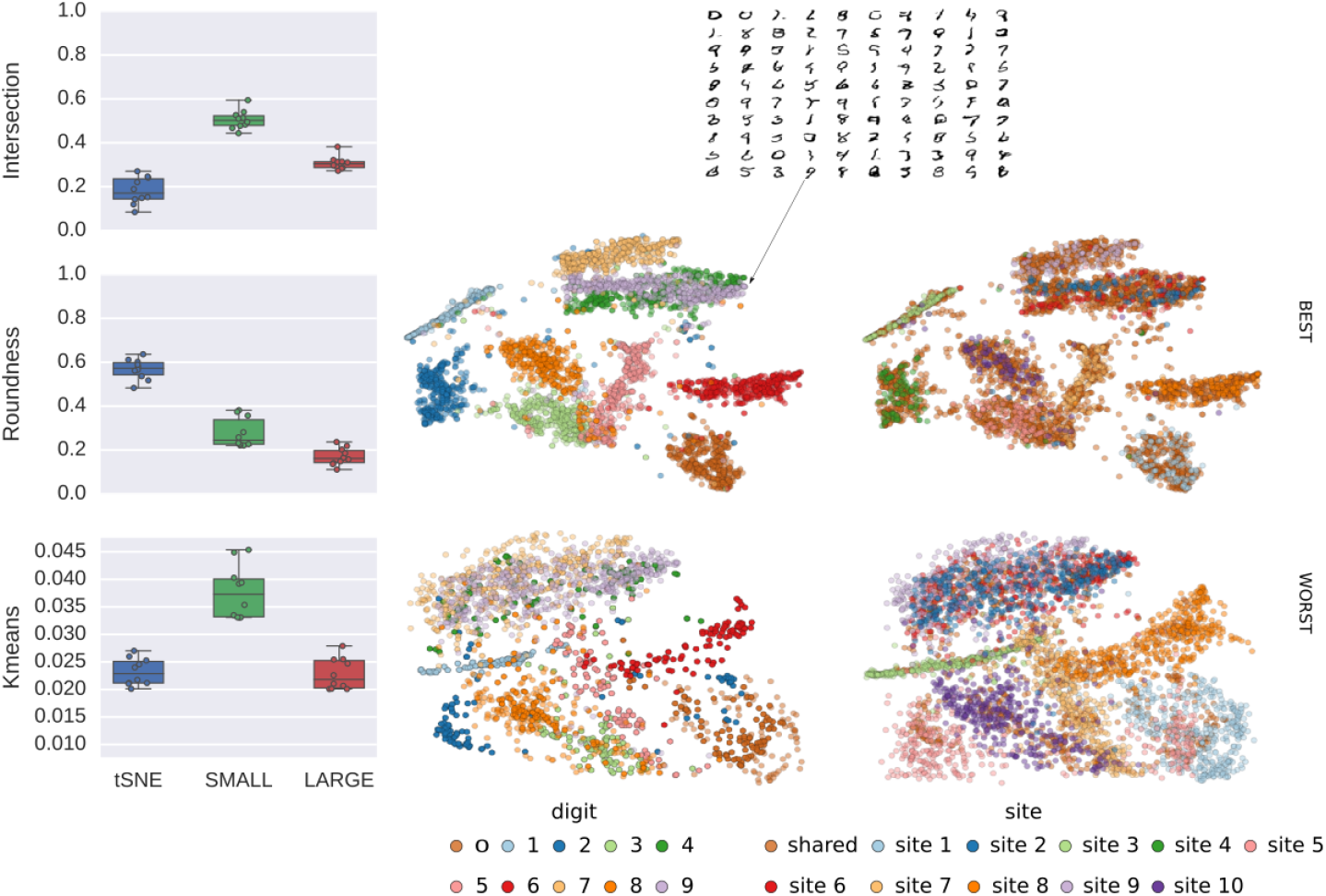
MNIST Experiment 1: Reference data contains samples of all of the MNIST digits, but is either a small or large amount. In the boxplots, tSNE was computed on pooled data and SMALL and LARGE represent smaller and larger reference datasets, respectively. Each row of the plots correspond to the best and worst performing runs. The left plots correspond to clusters labeled by digits, whereas the right plots correspond to clusters labeled by sites.

In Figure 2, the plots in each row correspond to the best and worst performing runs, respectively. The first column layout is colored by the digits, representing different clusters. The second column has layouts colored corresponding to sites. The box plots on the left show performance metrics computed by the three metrics proposed. t-SNE was computed on pooled, centralized data and SMALL and LARGE represent smaller and larger sizes of the reference dataset in dSNE runs. The comparison metrics show that performance is generally better when the shared portion of the data contains a large amount of data. However, the cluster roundness degrades with the size of the sample in the shared data. Thus, we observe this tradeoff between the roundness of the cluster and general performance. Overall, however, we observe that the dSNE clusters are less “round” compared to centralized t-SNE.

#### 5.1.2. COIL-20 Data

Figure 3 depicts the results of Experiment 1 on the COIL-20 dataset. Similar to that of Figure 2, the rows of the plots correspond best and worst dSNE runs. The first column has plots colored by each object, whereas the second column has clusters colored by sites. In the boxplots, t-SNE was computed on pooled data and SMALL and LARGE represent smaller and larger sizes of the reference datasets, respectively. The comparison metrics show similar results as the MNIST experiments. For large amounts of shared data, the comparison metric shows better performance than in the case of smaller samples in the reference dataset.

**Figure 3:**
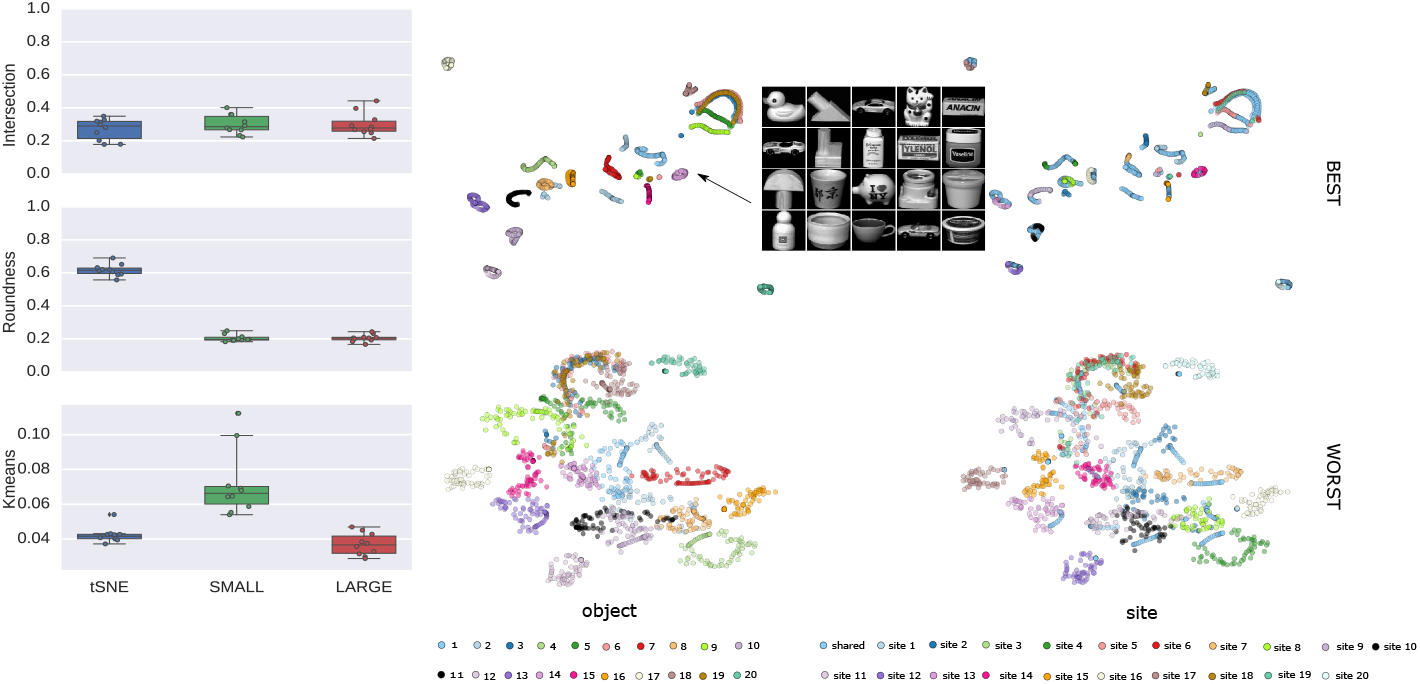
COIL-20 Experiment 1: reference data contains samples of all COIL-20 objects but is either in small or large amounts. In the boxplots, tSNE was computed on pooled data and SMALL and LARGE represent smaller and larger reference datasets, respectively. Each row of the plots correspond to the best and worst performing runs. The left plots correspond to clusters labeled by digits, whereas the right plots correspond to clusters labeled by sites. The COIL-20 dataset consist of 20 different objects which is shown in the figure. In the layout each point represents a object from these 20 objects.

Figure 4 depicts the results of Experiment 2 of the COIL-20 dataset. The comparison metrics show that we always obtain better results when the reference sample size is larger. For smaller reference samples, we observe highly overlapped clusters for which it is hard to distinguish the clusters for different objects.

**Figure 4:**
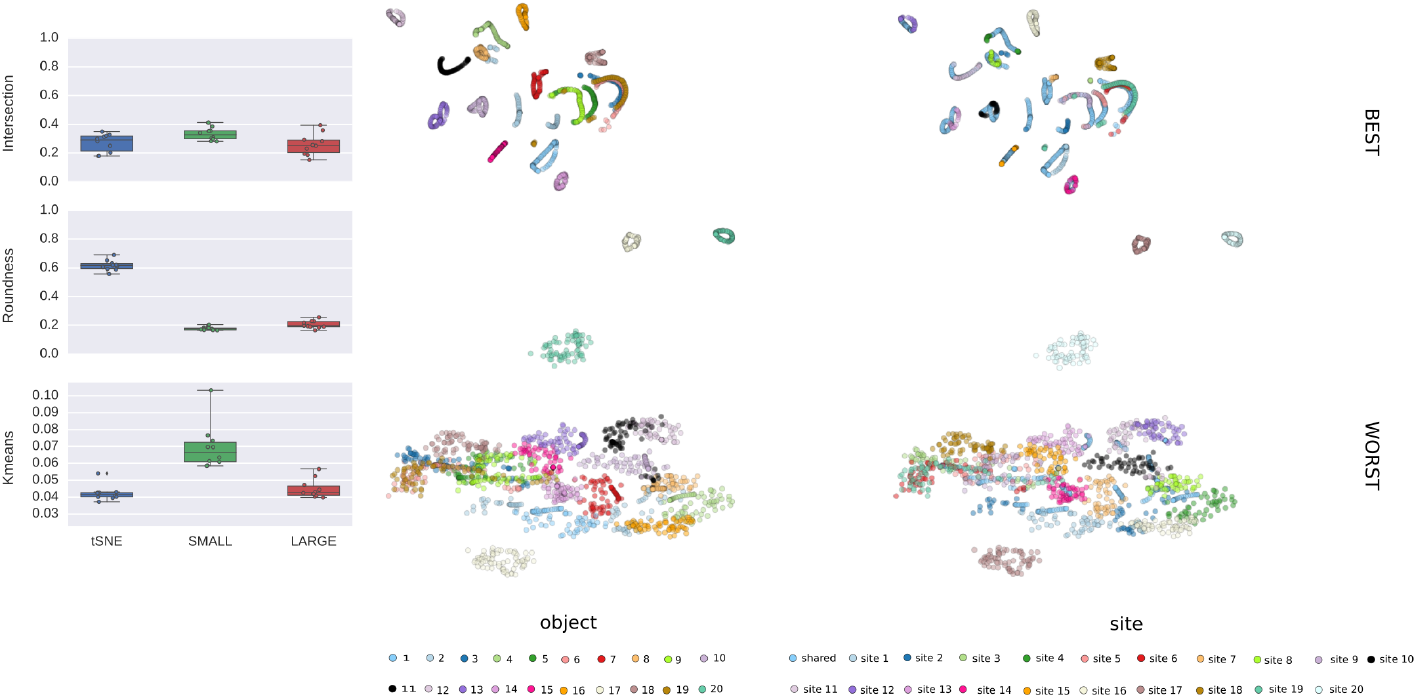
COIL-20 Experiment 2: the reference dataset is missing one unique COIL-20 object that is present at one of the local sites. In the boxplots, tSNE was computed on pooled data and SMALL and LARGE represent smaller and larger reference datasets, respectively. Each row of the plots correspond to the best and worst performing runs. The left plots correspond to clusters labeled by digits, whereas the right plots correspond to clusters labeled by sites.

### 5.2. Comparison of t-SNE and dSNE on Biomedical Data

We investigate the performance of multi-shot dSNE in comparison with the embedding produced by t-SNE on the pooled data using the QC metrics of the ABIDE fMRI, sMRI, PING, fBIRN, BSNIP and fMRI datasets.

#### 5.2.1. ABIDE fMRI Data

The layout of dSNE from the ABIDE dataset is shown in Figure 5. In every experiment, a total of 10 local and one remote sites are participating in the computation. The result from a centralized t-SNE run shows 10 different clusters. For each of the three random seed experiments of our decentralized simulation, we obtain 10 different clusters as well. In the layout, the samples that belong to the same site are marked the same color. In experiment 1, samples from site 1 and 4 are homogeneous. That’s why they are grouped together in the final embedding. In each of the experiments, we observe good separation among the clusters and consistent grouping of homogeneous samples, which shows that dSNE produces stable embeddings.

**Figure 5:**
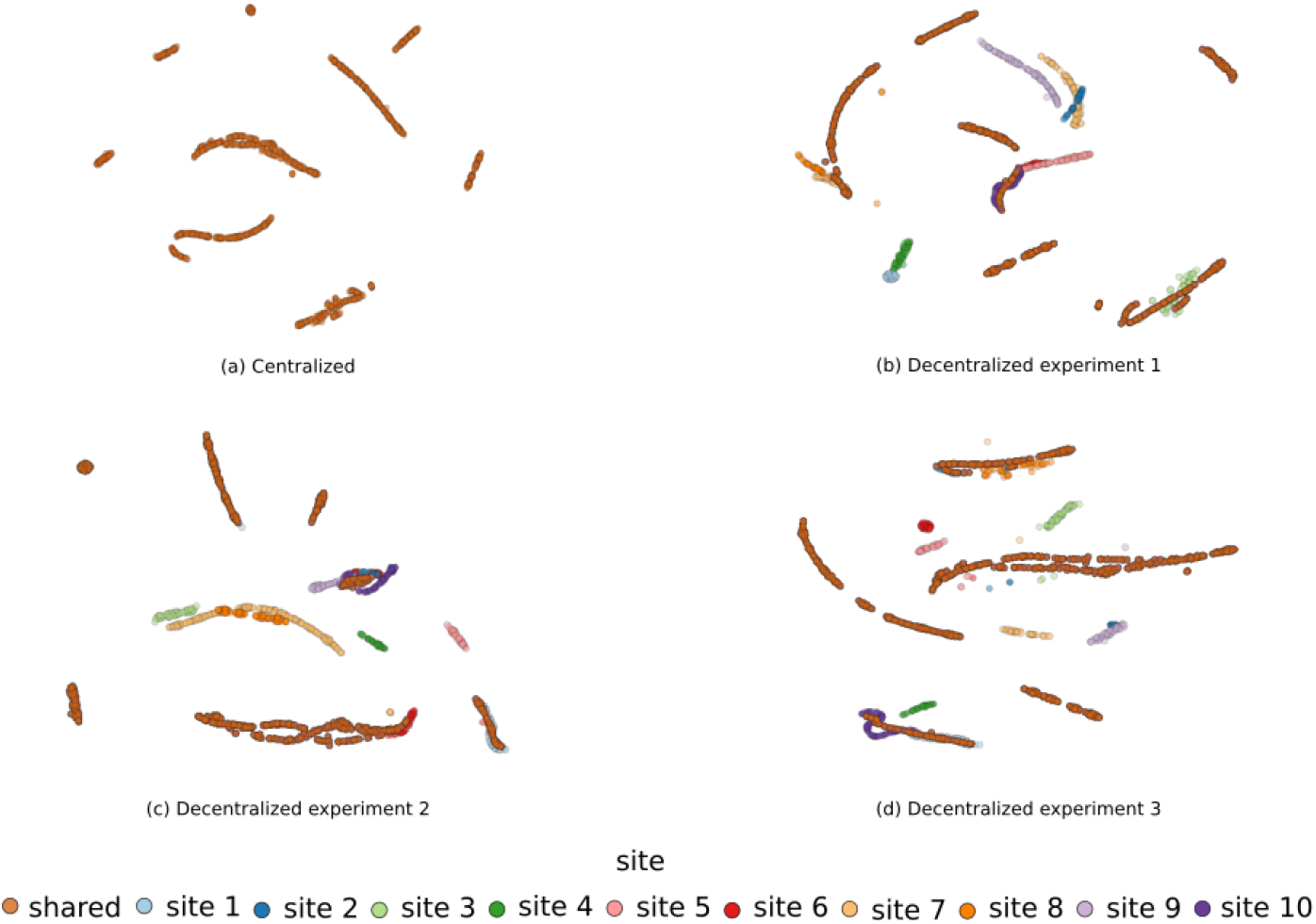
Experiment for the QC metrics of the ABIDE dataset. (a) the tSNE layout of pooled data; (b),(c) and (d) are the dSNE layouts for the three different experiments. In each dSNE experiment, 10 local and a coordinator sites participate in the computation. Similar to tSNE, in the decentralized setup, we get ten clusters, where each site is marked by unique color.

#### 5.2.2. sMRI Data

Figure 6 depicts embeddings produced by dSNE on sMRI data in our three experiments specified in Section 4.1.7. In Experiment 2, we obtain poor results, where all clusters overlap. An unrepresentative reference dataset predictably leads to such a layout. However, in Experiments 1 and 3, results closely resemble those obtained by tSNE. Note, that in all successful experiments scans of children younger that 11 formed a distinct separate cluster. Meanwhile the other age groups, although connected together in a single contiguous cluster, are ordered according to the age. Although the categories are discrete, it may be worth further investigation to inspect whether the age transition is smooth. This is an example of exploratory data analysis, where the structure of the resulting embedding may reveal some inherent regularities in a dataset. The age groups may not be as interesting, but serve as a clear demonstration that it is possible to use dSNE for discovery of data properties not already known to the researcher, similar to [52]. The dSNE can be used as visual and thus quick, intuitive and interpretable quality assurance, outlier detection, assessment of compatibility of datasets, and even assessment of site effects.

**Figure 6:**
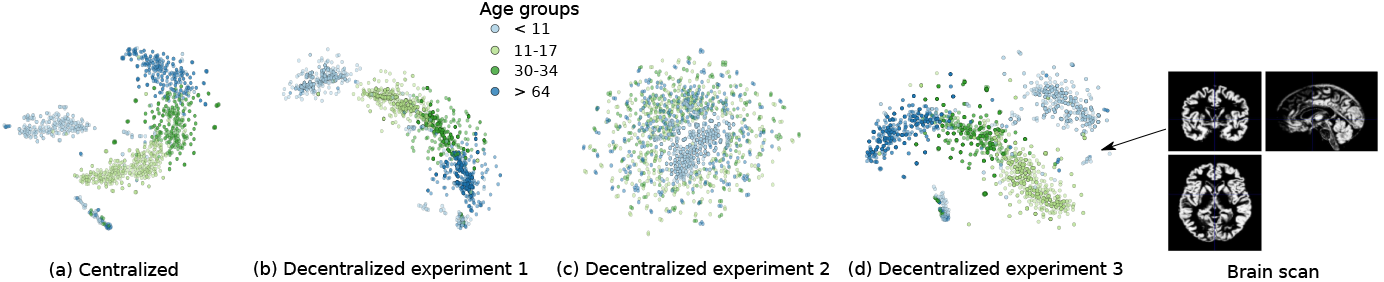
Experiment for the QC metrics of the sMRI dataset. (a) t-SNE layout of pooled data. (b), (c) and (d) are the dSNE layouts for the three different experiments. In all of the experiments, there are four total classes corresponding to an age group each and each class is marked by a unique color. The sMRI dataset consists of brain scans from different age group people and one of the brain scans is shown in the figure. These scans are preprocessed before entering the dSNE algorithm. In the layout, each point represents a single individual.

#### 5.2.3. PING Data

Figure 7 depicts the experimental results of the PING dataset. In both centralized t-SNE and dSNE, we get four major clusters. The fact that the number of clusters in the pooled scenario is equivalent to the number of clusters in the decentralized case shows promising results.

**Figure 7:**
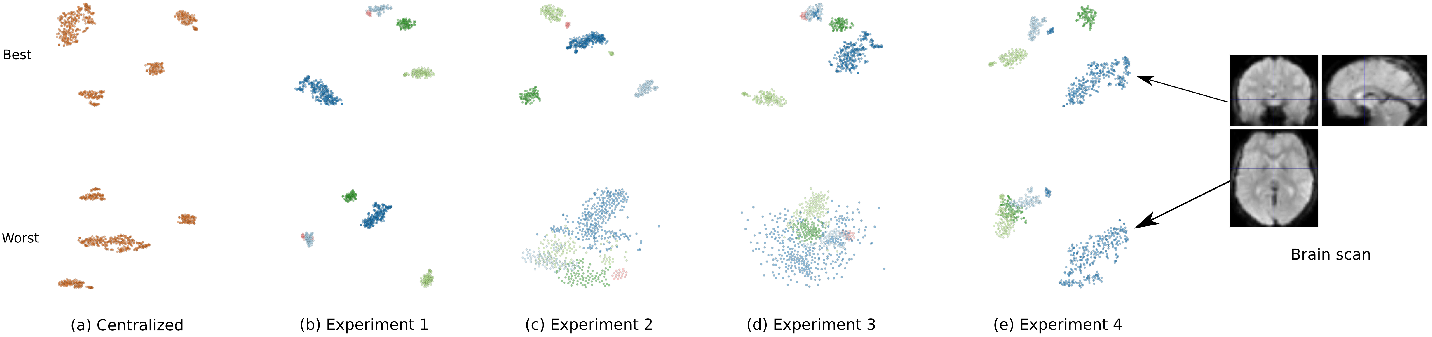
Experiment for QC metrics of the PING dataset. (a) the tSNE layout of the pooled data. In all of the columns, the top figure presents the best performing run, while the lower one represents the worst performing run. (b), (c), (d) and (e) are the dSNE layouts for four different dSNE experiments. In all of the dSNE experiments, we get four clusters, just like the t-SNE case. The PING dataset consist of brain imaging data of children and adolescents and one of the scans is shown in the figure. These scan data are preprocessed first and give input to our algorithm. In the layout, each point represents a single individual.

#### 5.2.4. fBIRN and BSNIP Data

Figure 8 depicts the results of the fBIRN and BSNIP experiments. In the first column, we present the t-SNE layout, where the top and bottom plots correspond to fBIRN and BSNIP, respectively. We obtain relatively well separated groups (HC and SZ) for fBIRN but not for BSNIP. In the second column, we present the t-SNE layout of combined datasets (fBIRN and BSNIP), where the top and bottom figures are colored by groups and sites, respectively. We ran t-SNE on the combined datasets, but for the plots, we only show fBIRN subjects. This is because our objective is to see how subjects from the same group of the fBIRN dataset from different sites embed together in lower dimensional space. In the third column, we present the dSNE layout of the combined datasets, where the top and bottom figures are colored by groups and sites, respectively. Again, we only plotted fBIRN after running dSNE on the combined datasets. From the layout of both t-SNE and dSNE on the combined datasets, we notice that subjects from the HC group are densely clustered in both plots. We obtain less dense clusters for the SZ group, but get embeddings that are similar in both t-SNE and dSNE plots. However, in the dSNE layout, we get more overlaps between clusters compared to t-SNE. We believe this is a reasonable tradeoff, as there is no direct communication between the private local sites.

**Figure 8:**
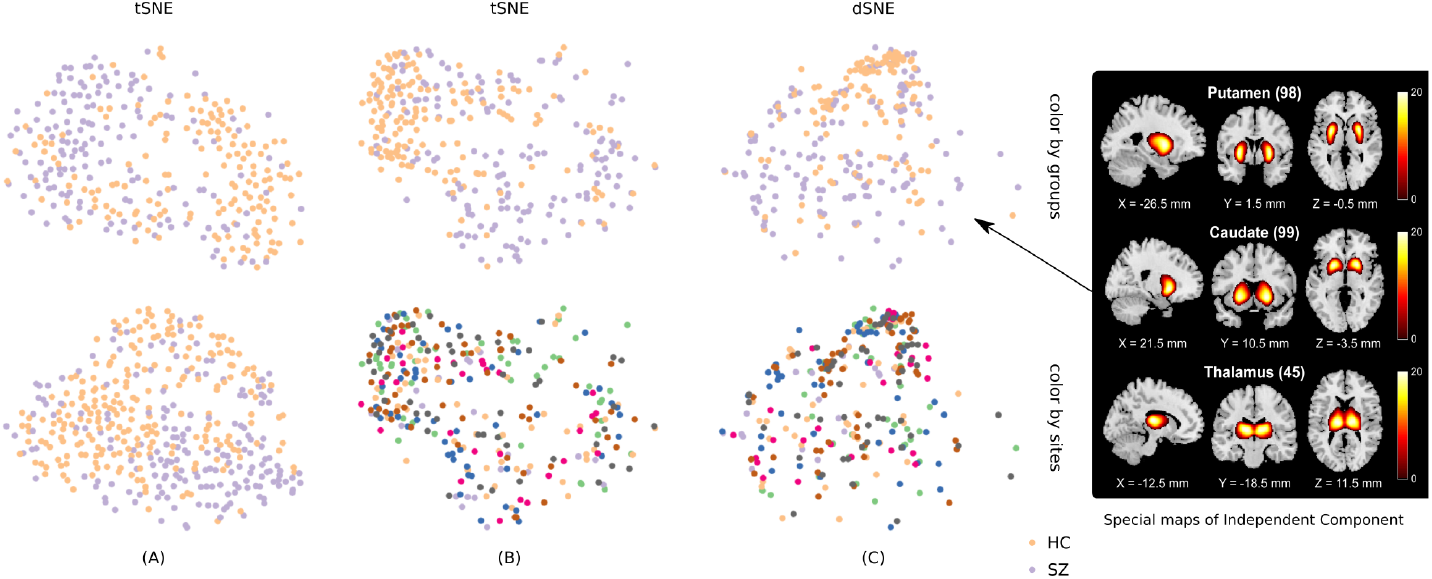
Experiment for the QC metrics of fBIRN and BSNIP datasets. (A) Top and bottom plots represent the t-SNE layout of fBIRN and BSNIP, respectively. (B) Top and bottom plots represent t-SNE layouts of combined (fBIRN + BSNIP) datasets but colored by groups and sites, respectively. (C) Top and bottom plots represent dSNE layouts of combined (fBIRN + BSNIP) datasets but colored by groups and sites, respectively. The fBIRN and BSNIP are the brain imaging data of healthy control and Schizophrenia. From these data intrinsic connectivity networks (ICNs) were extracted and used as input to our algorithm. One of the spatial maps of ICNs is shown in the figure.

It is also worth noting that during plotting, some points can be embedded in a very small dense region. To picture this better, we plotted the embeddings colored by sites. To our knowledge, we did not see any evidence of bias in which data are grouped by sites.

#### 5.2.5. fMRI Data

With the fMRI data, we demonstrate how we can perform quality control using t-SNE and dSNE visualizations. Figure 9 depicts the experimental results of fMRI data. In this figure, we observe four distinct clusters for both t-SNE on pooled data and dSNE on decentralized data. Having the same number of clusters in these cases highlight the potential of performing decentralized visualization with dSNE. Significantly, we can also observe the poor quality scan samples marked by the red cluster in the layout. The size of the red clusters in both plots are small and separated from the other samples. This shows that these samples are “outliers” from the other samples in the dataset.

**Figure 9:**
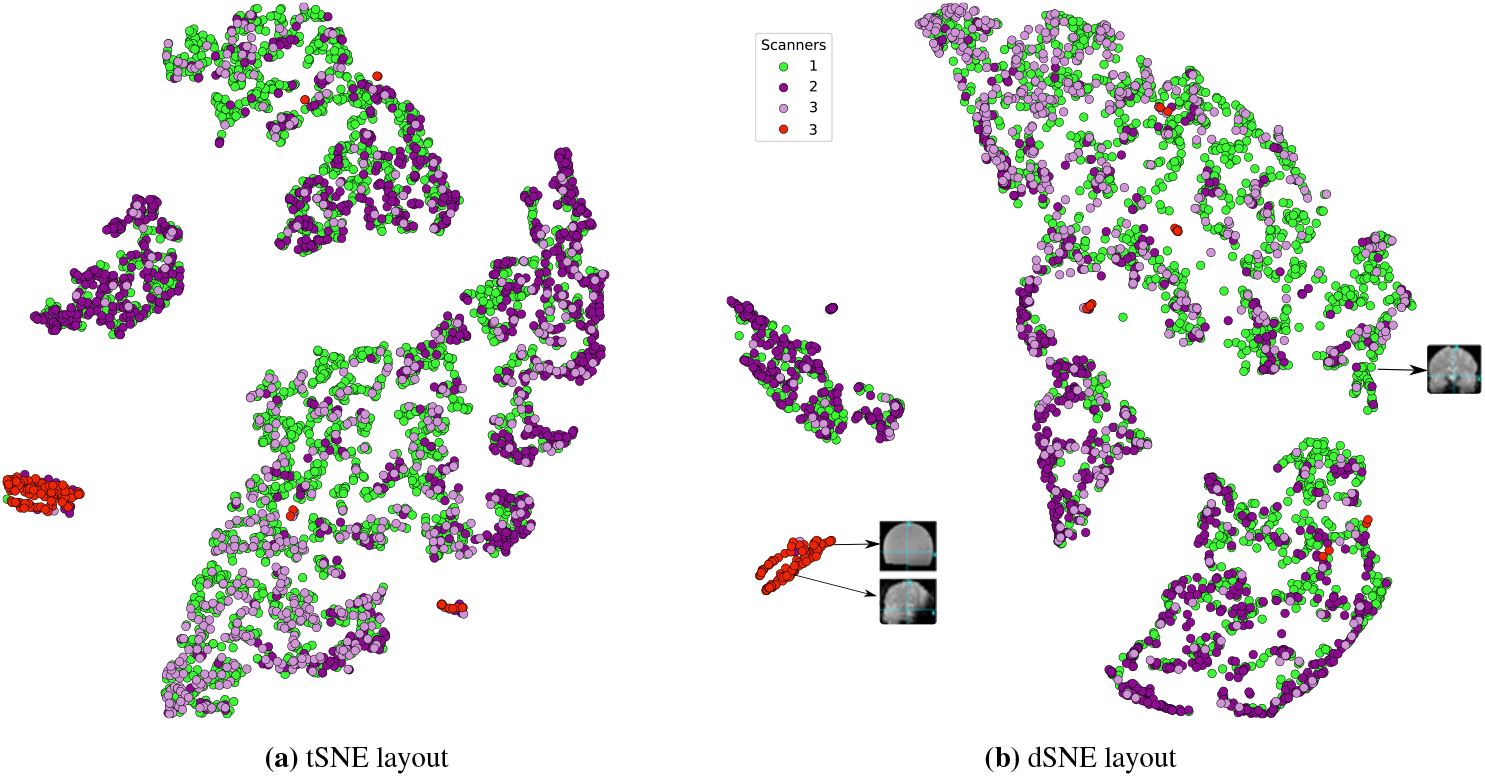
Experiment for the QC metrics of the fMRI dataset. Figures 9a and 9b are the layouts colored by Scanners for t-SNE and dSNE, respectively. In both experiments, we get four distinct clusters. Here, we can identify poor quality scan samples marked by the red cluster. In this experiment, three local and one remote sites participated in the computation. In the layout, each point represents a brain scan of an individual.

### 5.3. Comparison Between t-SNE and DP-dSNE

Lastly, we make a comparison between t-SNE, dSNE, and DP-dSNE on the MNIST and PING datasets. Figure 10 depicts the experimental results of our differentially private algorithm after 1000 iterations. In this figure, the left, middle, and right columns correspond to centralized t-SNE, dSNE and DP-dSNE, respectively. For all experiments, and in all plots, we see four different clusters corresponding to each class. Each cluster is distinct and well-separated amongst the other clusters. This proves to show the potential of our DP-dSNE algorithm, in which we can provide high utility whilst providing privacy of sensitive datasets in distributed locations.

**Figure 10:**
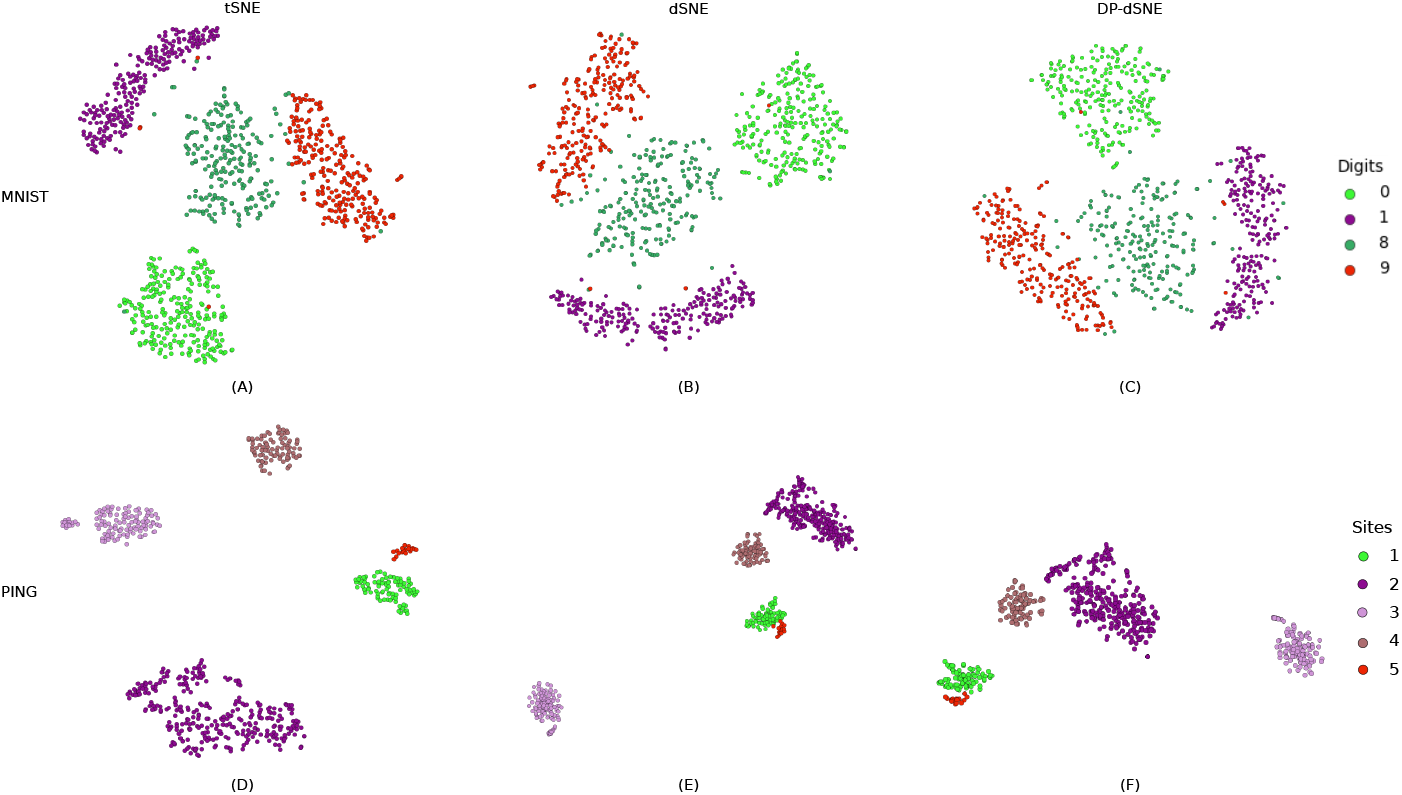
Experiment for DP-dSNE of MNIST and PING dataset with *σ*^2^ = 0.001. (A), (B), and (C) are the t-SNE, dSNE, and DP-dSNE layout for the MNIST dataset; (D), (E), and (F) are the t-SNE, dSNE, and DP-dSNE output for the PING dataset, respectively. We observe that DP-dSNE gives overall close results to dsne and centralized t-SNE. In the MNIST layout, each class is marked by a unique color and in PING layout, each site is marked by a unique color.

Figure 11 shows a plot of *ϵ* for a fixed *δ* for the PING dataset as we increase the number of iterations for convergence. Again, since each site has disjoint data samples from the other sites, we only observe *b*_max_ of the first site to compute *ϵ*. We also compare the (*ϵ*, *δ*) pairs of the RDP analysis compared to the strong composition [53]. Both the moments accountant and the RDP method are shown to give tighter, stronger privacy bounds [29, 36]. After 1000 iterations, for the PING dataset, given *δ* = 10^−5^ and *σ*^2^ = 0.001, we obtain (1.39,10^−5^)-DP using the moments accountant and RDP method and (5.85,10^−5^)-DP using strong composition. These (*ϵ*, *δ*) pairs are similar for the MNIST dataset.

**Figure 11:**
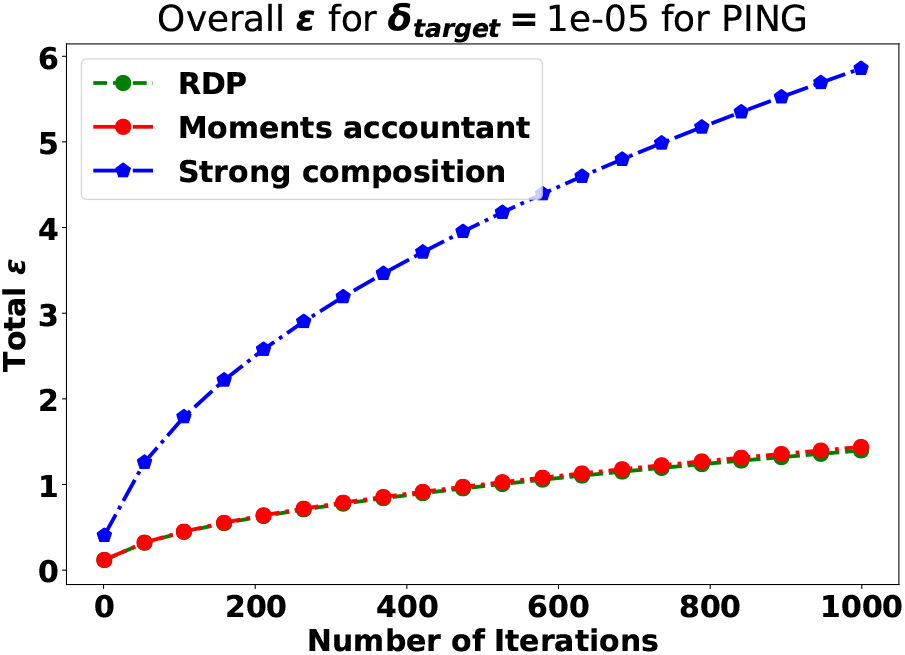
Plot of the number of iterations (*J*) vs. the total *ϵ* given *δ* = 10^−5^ and *σ*^2^ = 0.001. The RDP and moments accountant gives smaller values of *ϵ* over the strong composition method.

## 6. Discussion

The current practice of data sharing and pooling face great challenges as privacy concerns such as subject deidentification becomes more apparent. Previous studies have showed that in some cases it is possible to identify specific subjects from a dataset consisting of patients with rare diseases [54, 55]. The inability to combine datasets from different research groups and data sites can be devastating, as individual sites rarely contain enough data to answer the questions of foremost importance in biomedical research. There have been many notable methods that address the problem of data scarcity at individual sites in decentralized settings. For example, Virtual Pooling and Analysis of Research Data (ViPAR) [7] is a framework proposed in which a secure and trusted coordinator node (or server) synchronizes with the remote sites involved in the computation. At each iteration, each remote site sends data via an encrypted channel to the secure server. Data are then stored in RAM on the ViPAR coordinator server, where the data samples are analyzed and subsequently removed without ever being permanently stored. However, this process still relies on sending data outside of the original site. In addition, sending data via an encrypted channel incurs severe bandwidth and traffic overhead that ultimately increases computational load. Another promising approach is the Enhancing Neuroimaging Genetics through Meta-Analysis (ENIGMA) consortium [2], which is a community approach that requires the local sites to upload or email the summary statistics of the data following implementation of shared analysis scripts. ENIGMA uses both mega (if data pooling is possible) and meta analysis. In meta analysis, each local site runs the same analysis (e.g. regression) using the same measurements of the brain to aggregate summary statistics from all of the sites. The ENIGMA model has been widely embraced by the community. To run a meta analysis, the leading site has to coordinate with all of the local sites before starting and after the completion of computation. Meanwhile, the ENIGMA meta analysis approach does not support multi-variate or multi-shot computation, i.e. computing results in an iterative manner [56]. This is key as close to 50 percent of the data in some work groups, based on internal polling, cannot be centralized and thus much rely on meta-analysis. In addition, a standard meta analysis approach does not provide any formal guarantees that it will prevent the re-identification of individuals. In many machine learning problems, there are many cases in which statistics exchange must be done in a multi-shot manner, as single-shot is not enough to obtain an optimal solution [28].

The recent ongoing research on federated learning, differential privacy and encrypted computing is described in [57]. The Intel corporation has started a collaboration with the University of Pennsylvania and 19 other institutions to advance real world medical research using federated learning. Their work showed that a deep learning model trained by the traditional federated learning approach can reach up to 99% training accuracy [58].

Several notable tools and algorithms were introduced to handle federated computing efficiently. PySyft [59] is one of OpenMined’s Python code libraries that integrated cryptography and distributed technology with PyTorch and Tensorflow. This was mainly developed to train AI models in a secured way by ensuring patient privacy using distributed data. Our platform COINSTAC [4] is another example of an open source platform addressing these tasks. Researchers at Google Inc. introduced a model of federated learning using distributed data of user’s mobile devices [60]. In this model, a mobile device downloads and trains the model by accessing the data of the user’s device. It summarizes the changes and sends them as an update to the cloud using encrypted communication. Finally, the updates coming from all of the devices are averaged in the cloud and improves the shared model.

Federated Averaging (FedAvg) is a computation technique introduced in 2016 to fit a global model in the decentralized setting [60]. In this model, the parameters are initialized on the server and distributed to the local clients. After training the model on each local dataset over multiple iterations, the trained parameters are delivered to the server, which computes the average to send the weights back to the local sites. We adopted a very similar approach to Federated Averaging in dSNE [28] before proposing the proxy data sharing technique [61]. In dSNE, each local site accesses a publicly available dataset and updates its model using the combination of the shared data and its own local data. Similar to the communication round in [61], each site runs the operations over a fixed number of iterations to reach the optimal solution. We also applied the averaging technique in which the local model is averaged after each iteration and transferred to by the coordinator node.

None of these existing methods fully address how we can perform data assessment and quality control in a decentralized manner. Data centers and institutions may not be willing or able to share their data due to a need to preserve the privacy of their subjects, precluding analyses that pool the data to a single site. To address these issues, we introduced a way to visualize federated datasets in a single display: dSNE [28] and its differentially private counterpart, DP-dSNE. In both algorithms, one coordinator node communicates with all local sites during one computation period. Our multi-shot approach follows from the averaging strategy similar to that of [56]. The performances of t-SNE and dSNE are presented in Figures 2–9. In the best case scenarios, dSNE almost replicates t-SNE and shows great performance in terms of the comparison metrics. We showed that the performance increases when the reference data contains a large amount of samples as shown in Figure 2 for the MNIST dataset. We observe the similar type of behavior for the COIL-20 dataset, shown in Figure 3 and Figure 4. Our results in the influence of large reference samples reflect the results also shown in djICA [62].

In Figures 5–9, we were also able to observe significant results for six different biomedical datasets (ABIDE, sMRI, PING, fBIRN, BSNIP, and fMRI). Data assessment and quality control plays a vital role during data acquisition from multiple data sources, especially to keep consistency or adjust parameters across various studies. We designed Experiment 4.7 to check the effectiveness of our algorithm in detecting outliers in multi-site consortia. In Experiment 4.7, we collected 705 samples from Boulder site, where 120 of the samples were acquired from the second 3T dataset that was from a specific study with a very different acquisition protocol. Most importantly, these 120 samples were examples of a poor quality scan. In Figure 9, we see that these specific samples are clustered together (marked by red color) as outliers in both t-SNE and dSNE plots. Our results are similar to one of our earlier works, which used t-SNE to detect outliers of the same fMRI dataset [10].

In Figures 10 and 11, we can see the performance of DP-dSNE with a noise variance of *σ*^2^ = 0.001 on the MNIST and PING datasets. From Figure 10, we see that DP-dSNE gives very similar results compared to t-SNE and dSNE. We implemented DP-dSNE as dSNE does not provide any formal privacy guarantees. Even though the data samples in each local site never leaves the sites, sending the gradient values from each local site could potentially leak privacy. For datasets that are sensitive and have stricter protocols, we propose that DP-dSNE can provide utility similar to that of dSNE with strong privacy guarantees.

Lastly, we also implemented single-shot dSNE, but the results were not as promising as multi-shot dSNE (dSNE), as shown in A.13. In single-shot dSNE, there is no way for the local sites to communicate iteratively. Additionally, a major problem of averaging arises when different sites are widely varied in terms of size and population [4]. This problem is analogous to the size of the reference dataset in multi-shot dSNE. The reference dataset should provide variability and be large in size, in which otherwise, there is a high chance of not obtaining the optimal embedding. To obtain an optimal embedding, the algorithm should be ran multiple times with a different initialization.

The Collaborative Informatics and Neuroimaging Suite Toolkit for Anonymous Computation (COINSTAC^6^) [4, 63], a dynamic and decentralized platform, was introduced to address the difficulty of data sharing. This platform gives scope to perform distributed computation by using the commonly used algorithm in privacy-preserving mode.

Our dSNE algorithm is currently deployed within the COINSTAC framework^7^. Figure 12 shows the different computation phases of dSNE experiment in COINSTAC using MNIST dataset. In this experiment, three local and one remote site participate in the computation, where the remote site contains 200 samples of each digit (0 to 9) and each local site contains 20 samples of each digit. The results shown in the Figure 12 demonstrates the robust capability of the COINSTAC framework. Future work consists of running simulations using all of the datasets used in this paper. This serves to demonstrate that COINSTAC ensures the real time applicability of our algorithm in the biomedical domain.

**Figure 12:**
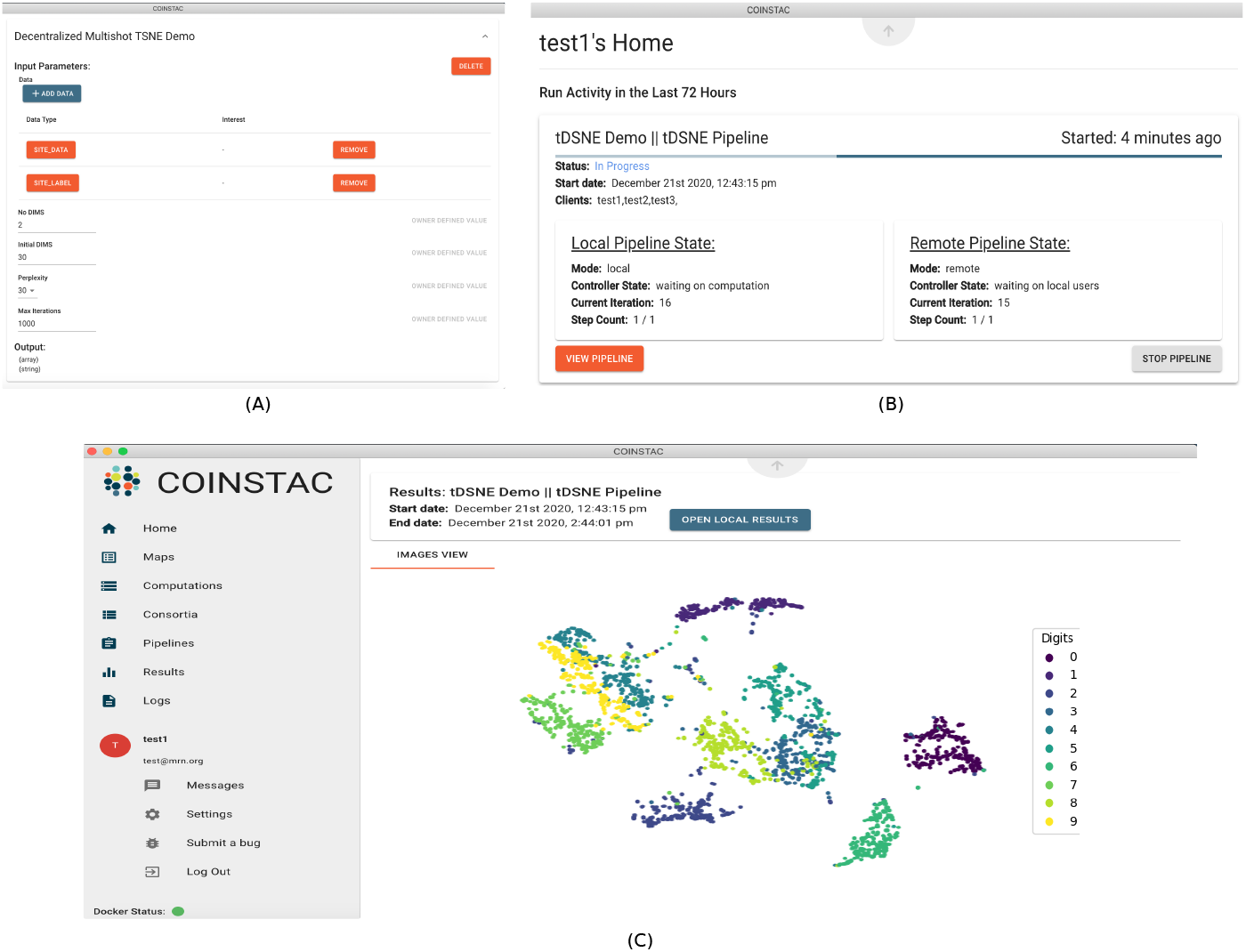
An run time demo of dSNE algoroithm in coinstac simulator. (A), (B) and (C) are the computation phase of dSNE at the beginning, middle, and at the end of the simulation

## 7. Conclusion

In this paper, we have proposed two algorithms: decentralized stochastic neighbor embedding (dSNE) and differentially private decentralized stochastic neighbor embedding (DP-dSNE). Our dSNE algorithm enables the embedding of high dimensional private neuroimaging data spread over multiple sites into low dimensional space for visualization. This visualization allows us to perform quality control of poor data samples and also helps learn a global interrelation structure among brain volumes or feature vectors. Throughout the dSNE computation, no data samples leave their respective local sites, and only minimal gradient information from the embedding space is transferred across the sites. The clusters in the output embeddings are formed by samples belonging to classes, possibly present across many locations. Of course, the algorithm is neither explicitly aware nor requires the prior existence of any classes. The main idea of this iterative method is to share only the parts related to the publicly available reference dataset. As our results show, this is enough to co-orient classes that are spread across multiple locations. Extensive validation of eight datasets (two toy and six multisite neuroimaging) demonstrate the utility of our approaches. Our results showed that although multi-shot dSNE is robust to various conditions and settings (e.g. changes in the number of sites, and rare or missing data), and highest performance is achieved when the reference dataset is dense. Even though the data samples never leave the local sites in dSNE, there is still room for a potential privacy leak, as we are sending the gradients over to a coordinator node. DP-dSNE tackles this by introducing formal privacy guarantees within the gradients. Our results show that both dSNE and DP-dSNE provide good utility for decentralized visualization whilst preserving privacy of the local samples. The implementation and integration of the algorithm with an existing neuroimaging platform for federated neuroimaging COINSTAC provides our methods as ready to use tools.

For future work, we believe that an alternative solution to our decentralized setting can be to use an average of the gradients weighted by the quality of the respective local t-SNE embeddings. However, it is not immediately clear how one should approach this. Using clustering measures on location specific data to weigh each *Y* may bias the results toward good local groupings over poor ones. Our novel metrics are also not quite able to convey each site’s contribution to give it proper weight. In most general settings, we do not know a priori what type of data samples each site contains, as each local site has private data. Given these difficulties, we leave the problem for future work, noting that it could be an exciting research direction. Finally, we conclude with that we believe dSNE is a valuable quality control tool for virtual consortia when working with private data in decentralized settings.

## Acknowledgement

This work was supported by NIH 1R01DA040487, R01DA049238, R01MH121246, 2R01EB006841, 2RF1MH121885.

Autism data was provided by ABIDE. We acknowledge primary support for the work by Adriana Di Martino provided by the (NIMH K23MH087770) and the Leon Levy Foundation and primary support for the work by Michael P. Milham and the INDI team was provided by gifts from Joseph P. Healy and the Stavros Niarchos Foundation to the Child Mind Institute, as well as by an NIMH award to MPM (NIMH R03MH096321).

Data for Schizophrenia classification used in this study were downloaded from the Function BIRN Data Repository (http://bdr.birncommunity.org:8080/BDR/), supported by grants to the Function BIRN (U24-RR021992) Testbed funded by the National Center for Research Resources at the National Institutes of Health, U.S.A. and from the COllaborative Informatics and Neuroimaging Suite Data Exchange tool (COINS; http://coins.trendscenter.org) and data collection was performed at the Mind Research Network, and funded by a Center of Biomedical Research Excellence (COBRE) grant 5P20RR021938/P20GM103472 from the NIH to Dr.Vince Calhoun.

## Appendix A. Single-shot dSNE

For single shot d-SNE (Algorithm 8), we first pass the reference data from centralized site *C* to each local site.

Now each local site has data consisting of two portions: (1) its local dataset, for which we need to preserve privacy, and (2) the shared reference dataset. Each local site runs the t-SNE algorithm on this combined data (local and reference) and produces an embedding into a low dimensional space. However, while computing each iteration of tSNE, a local site computes gradient based on combined data, but it only updates the embedding vectors **y** for the local dataset. The embedding for the shared data has been pre-computed at the coordinator node and shared with each local site. Similar to the landmark points approach [26], our method uses reference points to tie together data from multiple sites. In practice, the samples in the shared dataset are not controlled by the researchers using our method, and it is hard to assess the usefulness of each sample in the shared data in advance. In the end, each local site obtains an embedding of its data and the embedding of the shared dataset. Since the embedding points of the shared dataset do not change, all local embeddings are easily combined by aligning the points representing the shared data.

**Algorithm 8.**
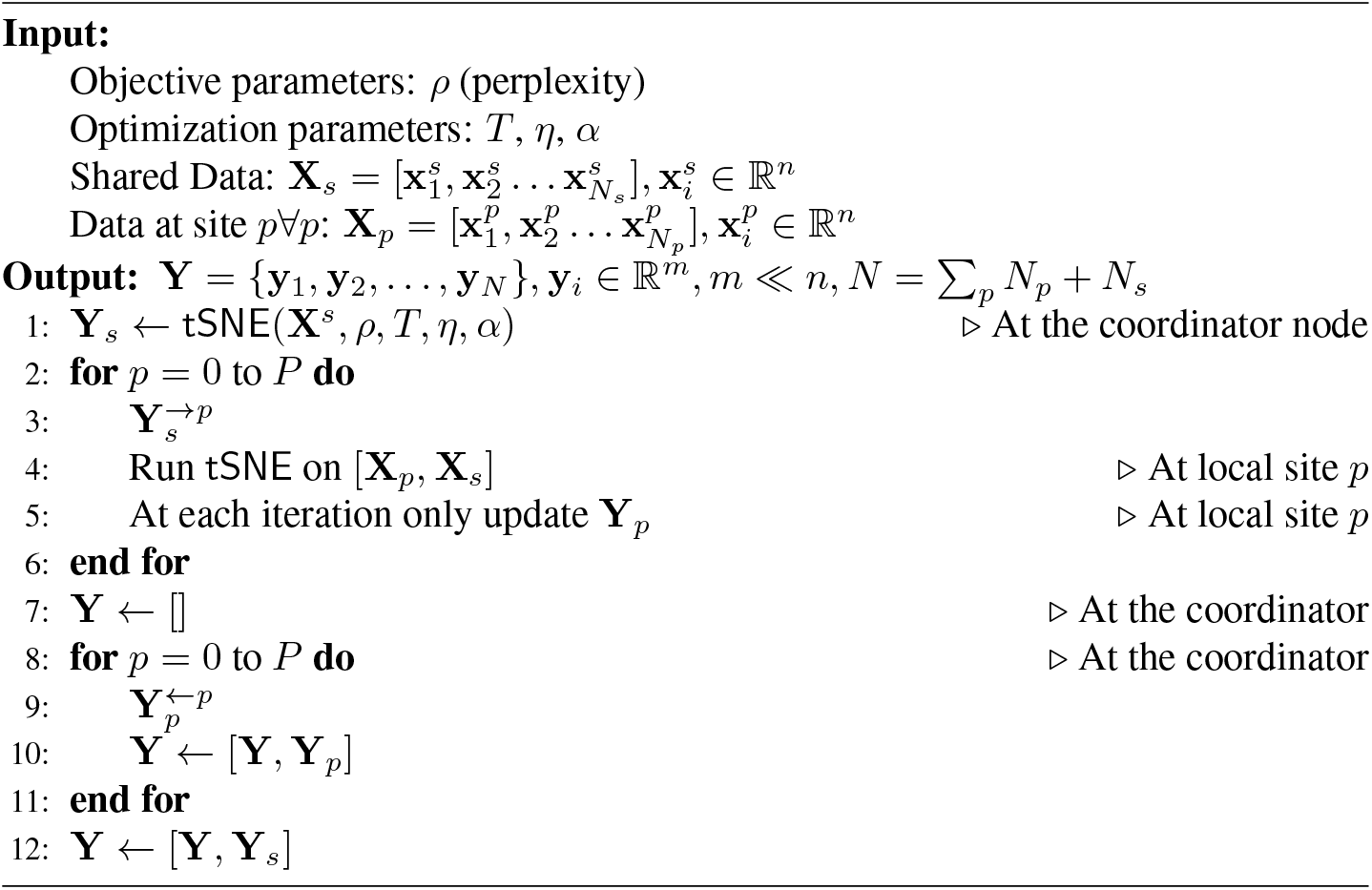
singleshotDSNE

**Figure A.13:**
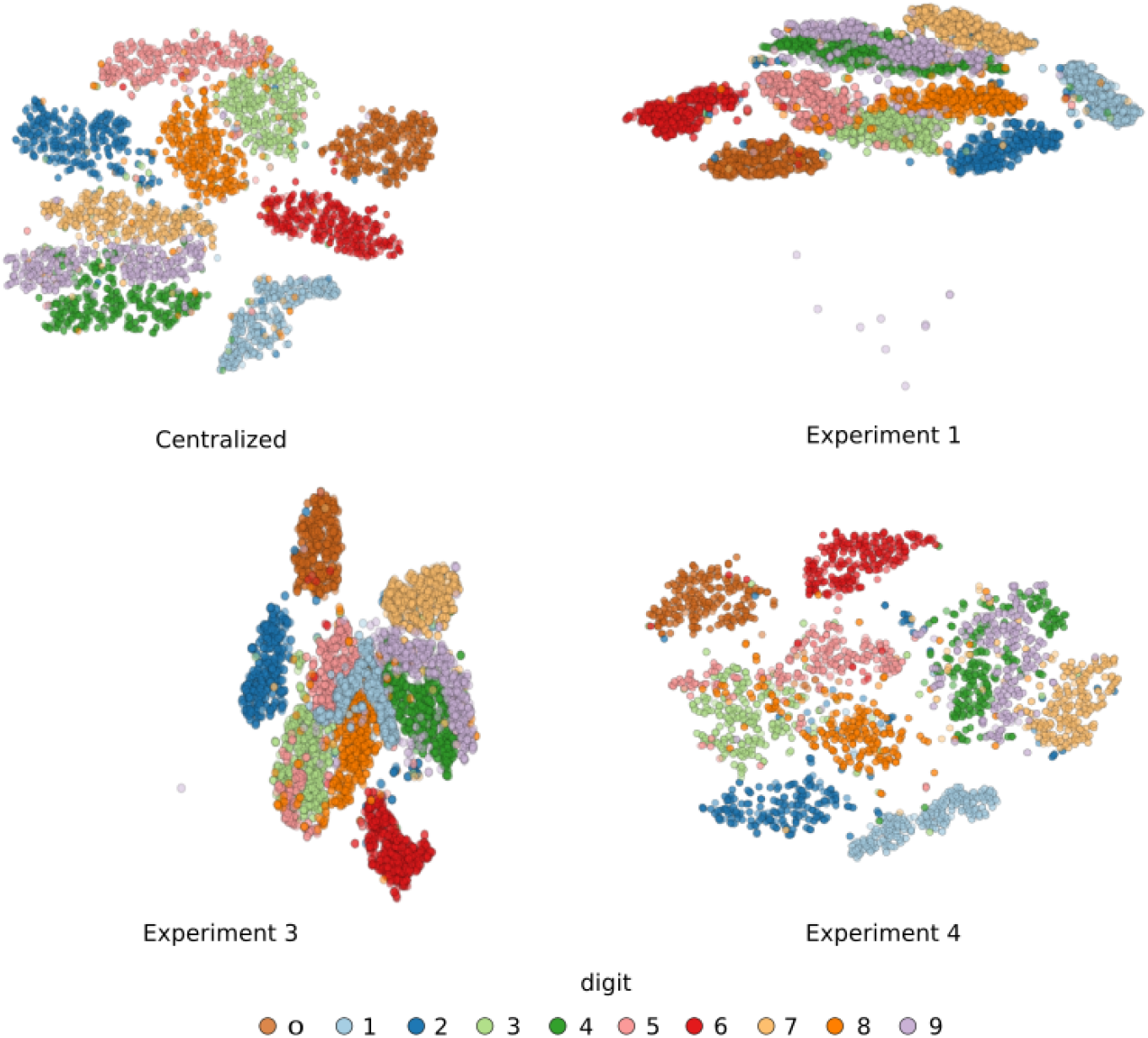
Single-shot dSNE layout of MNIST data. Single-shot was run for the experiment of 1, 3 and 4 of MNIST dataset. For all experiments, we are able to embed and group same digits from different sites with-out passing any site info to others. Here every digit is marked by a unique color. Centralized - is the original tSNE solution for locally grouped data. Digits are correctly grouped into clusters but these clusters tend to heavily overlapped.

## Footnotes

1 https://www.kaggle.com/c/digit-recognizer

2 http://www.cs.columbia.edu/CAVE/software/softlib/coil-20.php

3 http://fcon_1000.projects.nitrc.org/indi/abide/

4 http://pingstudy.ucsd.edu/Data.php

5 The full list is available here: https://github.com/preprocessed-connectomes-project/quality-assessment-protocol/tree/coordinator/normative_data

6 https://coinstac.org/

7 The code is available at: https://github.com/trendscenter/coinstac-dsne-multishot

## Notes

### Competing Interest Statement

The authors have declared no competing interest.

